# Information-Guided Planning for Online System Identification of Partially Observed Nonlinear Dynamical Systems

**DOI:** 10.64898/2026.10.05.756678

**Authors:** Hyungju Jeon, Il Memming Park

## Abstract

Online identification of nonlinear dynamical systems faces two challenges. First, noisy, partial observations make the latent state difficult to estimate. Second, local attractors can also keep passive trajectories in regions that reveal little about the unknown dynamics. Both challenges commonly arise in closed-loop neural experiments where stimulation must be chosen online from noisy recordings of neural activity that may be confined to a small part of the state space and produce highly redundant observations. To address this problem, we introduce PALDI (Planning-based Active Latent Dynamics Identification), which combines online system identification and state estimation with finite-horizon planning to actively select stimulations that maximize expected information gain. Our objective combines sensitivity of dynamics with state Fisher information, reweighted by latent-state uncertainty. We compare PALDI with several baselines on two multistable neural circuits: a neural mass model and a decision-making spiking network model. In the neural mass model, we show that PALDI remains robust under realistic recording conditions where observations have a low signal-to-noise ratio or are less informative in some regions of the state space. In the spiking network, we show that despite model mismatch, PALDI learns a model accurate enough to derive an efficient control policy, which overturns the circuit’s decision more reliably than the baselines under the same control budget. These results show that PALDI provides accurate, robust, and sample-efficient identification of neural circuit dynamics and suggest its potential as a practical strategy for learning models in closed-loop neural experiments.

## I. Introduction

**I**dentifying unknown parameters in a nonlinear dynamical system from observed time series is a central problem in control, signal processing, and adaptive modeling. In many practical settings, the system state is only partially observed, the dynamics are nonlinear, and the measurements are corrupted by noise that may be non-Gaussian. Moreover, in many applications, real-time constraints necessitate that inference and learning be performed online as new observations arrive, imposing computational and latency constraints [1]–[3].

Closed-loop neurotechnology is a particular example in neuroscience where these challenges arise. While simple threshold-crossing rules have been shown to improve therapeutic outcomes [4], [5] and to enable experimental control of neural activity [6], [7], more effective control requires an accurate model that can predict how the system evolves over time [8].

In practice, such a model is learned from passive recordings, which often cover only part of the state space. Attractors, common in biological and engineered systems, confine trajectories to a small region of the state space. Neural population activity, for example, is commonly modeled by recurrent dynamics that converge to fixed points, limit cycles, or low-dimensional manifolds associated with memory states, decisions, or motor primitives [9]–[12]. With multiple attractors or metastable dynamics, activity can stay in one basin for long periods and leave it only rarely. Passive recordings thus only weakly inform the dynamics in the unvisited regions [12]–[15]. A passively learned model that makes no explicit assumptions about the unobserved dynamics then has limited use for complex control that requires knowledge of the entire dynamical landscape.

This lack of information can be addressed through active system identification, which selects stimulation or inputs that make future observations informative about the unknown dynamics [16]–[19]. Appropriately chosen inputs can drive activity out of attractors to explore unvisited regions and reach those where observations are more informative. These methods, however, typically (1) assume a fully observed state, whereas neural experiments observe the state only through noisy recordings, and (2) lack planning over the several time steps needed to leave an attractor.

We develop PALDI (Planning-based Active Latent Dynamics Identification), which combines online approximate Bayesian state and parameter inference with finite-horizon planning. PALDI assumes a parametric latent dynamical model whose functional form is known and selects inputs that maximize the expected information gain about its parameters. We approximate this information gain with a computationally efficient objective that steers the system toward regions where (A) the state trajectory over the planning horizon is sensitive to the parameters (parameter sensitivity), (B) the (partial) observations are sensitive to the state (state Fisher information), and (C) the state estimate is precise (uncertainty discount).

In the following, we derive our proposed algorithm PALDI from first principles and evaluate it on two multistable neural circuits: a neural mass model [20] and a decision-making spiking network [21], where the true circuit lies outside the agent’s model class. We test how observation quality and individual components of the acquisition objective affect exploration and identification. We further evaluate the identified models through a control task, testing whether they can guide effective interventions.

## II. Methods

### A. Method Overview

Online active identification requires simultaneous filtering and planning. The filter must estimate an unobserved state and system parameters from noisy data, while the planner predicts the informativeness of future observations that depend on the current state and parameters. Effective active identification therefore requires both accurate state estimation and understanding of how system behavior depends on the parameters. PALDI runs the filter and the planner in a single loop (Figure 1). At each time step, an extended Kalman filter (EKF) updates the latent-state estimate from each new observation [1], [22]. After a block of steps, the parameter estimate is updated from the information accumulated in the block, following a dual state–parameter recursion [23]. Every few steps, the planner uses the improved cross-entropy method (iCEM) [24] to find the input sequence whose predicted observations would most reduce parameter uncertainty, an approximation to Bayesian expected information gain [25]–[27].

**Fig. 1.**
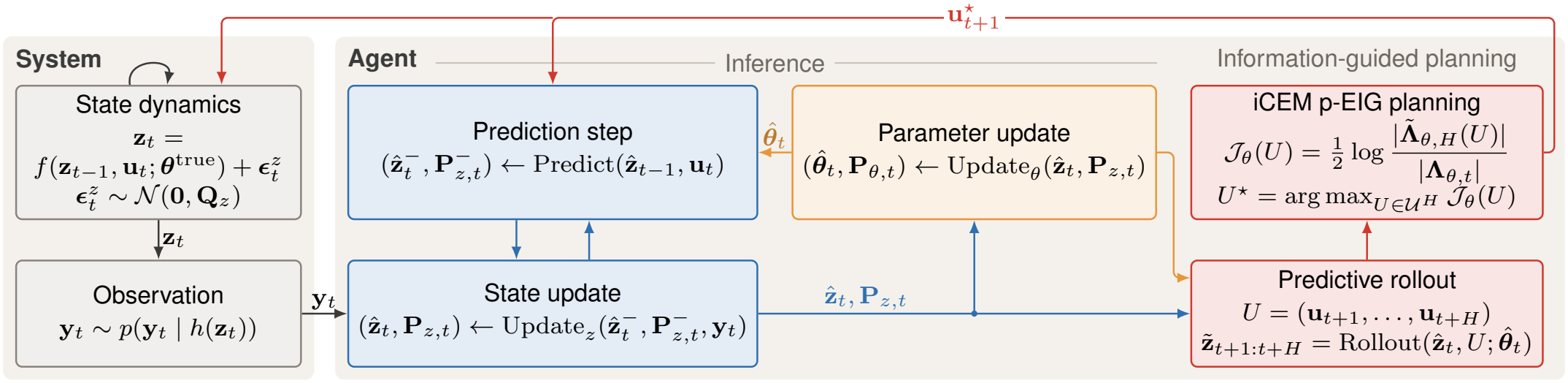
Information flow in the proposed framework. Observations update beliefs over latent states and parameters, and estimated information guides input selection. Blue and orange arrows carry the state and parameter beliefs; red arrows show planning and the selected input 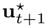.

**Fig. 2.**
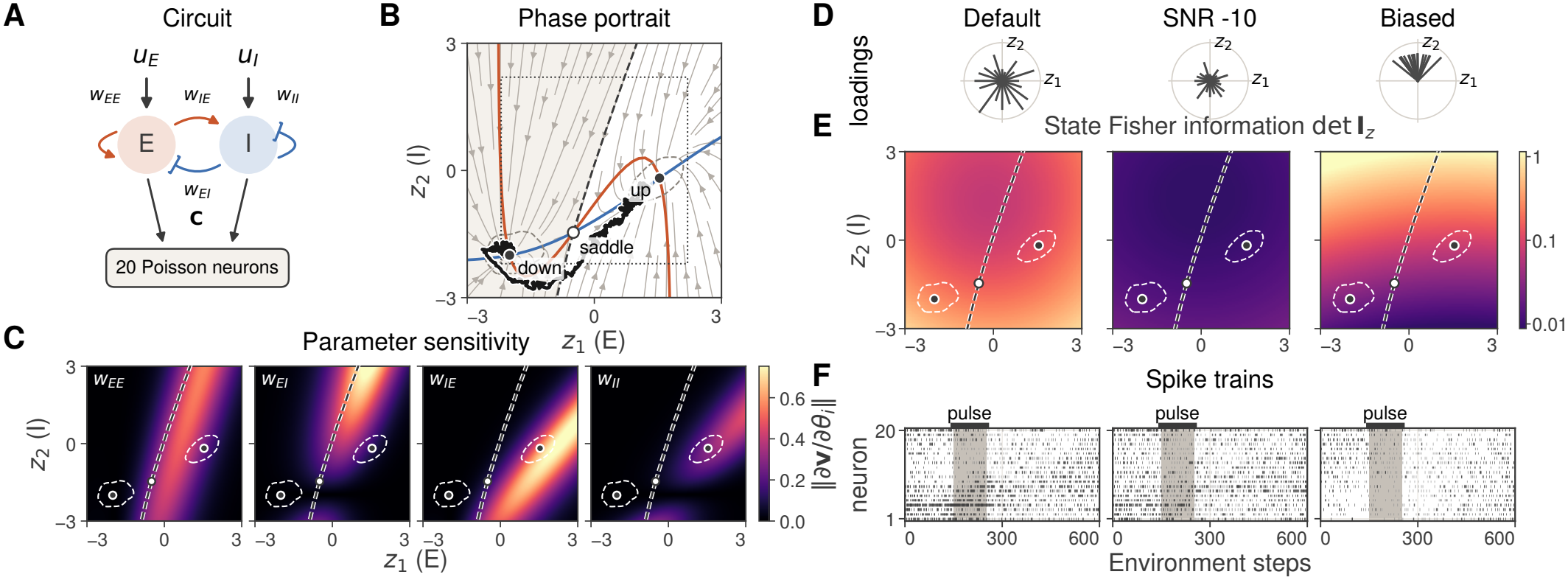
Neural mass model dynamics and observations. Model, parameter sensitivity, and observation models. (**A**) Circuit with four unknown weights, population-specific inputs, and a Poisson readout. (**B**) Input-free phase portrait: excitatory and inhibitory nullclines (orange, blue), stable states (filled), a saddle (open), and the separatrix (dashed) between the down-state (shaded) and up-state basins; the dotted square marks the evaluation region. (**C**) Parameter sensitivity ∥*∂***v***/∂θ*_*i*_∥. Dashed contours in (**B**,**C**,**E**) enclose 90% of passive states. (**D–F**) Observation models with uniform loading at *−*5 and *−*10 dB and biased loading at *−*6 dB. (**D**) Loading directions, the rows of **C**. (**E**) Determinant of the state Fisher information **I***z*. (**F**) Spike trains along the black path in (**B**), which an illustrative input pulse (gray) drives from the down to the up state.

### B. State-Space Model

We use a nonlinear state-space model [1], [22], [28]. The latent state 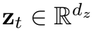 evolves as

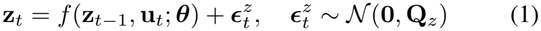

where *f*(·) is the discrete nonlinear transition map with unknown parameters 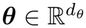, **u**_*t*_ is the known input applied between observations *t*−1 and t, and **Q**_*z*_ is the process-noise covariance per step. We write ***θ***^true^ for the parameters of the data-generating system and reserve the superscript ⋆ for optimized quantities.

Observations **y**_*t*_ ~p(**y**_*t*_| **z**_*t*_) depend on the latent state through a known observation model. Treating the observation model as known isolates identification of the latent dynamics: if both the observation model and f were unrestricted, a change in the observation model could be indistinguishable from a change in the latent coordinates or dynamics. Our primary interest is in neural recordings, where each observation **y**_*t*_ is a vector of binned spike counts from d_*y*_ neurons that gives partial and noisy access to the latent dynamics.

We therefore use the standard Poisson log-linear observation model [29],

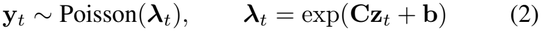

Each component of ***λ***_*t*_ is the expected spike count of one neuron in a time bin of width Δt seconds, so the firing rate in Hz is **r**_*t*_ = ***λ***_*t*_/Δt.

### C. Online Approximate Inference

Exact filtering for p(**z**_*t*_, ***θ*** |**y**_1:*t*_) is generally intractable for nonlinear dynamics with non-Gaussian observations. A full joint Gaussian approximation also requires a covariance in 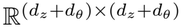. We instead use the dual-estimation structure [23], which maintains separate Gaussian beliefs over the latent state and parameters,

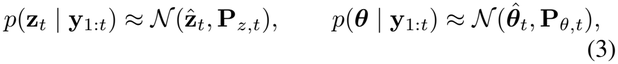

and update them in sequence. At each step, an EKF updates the latent state with the parameter estimate fixed. The resulting state estimates provide local score and Hessian increments for parameter learning.

#### 1) State Update

With parameters fixed at the current estimate 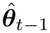, which changes only at parameter-update times (Section II-C2), the state prediction follows the standard EKF linearization [1], [22]:

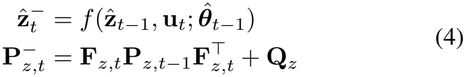

where **F**_*z,t*_ = ∂f/∂**z** is evaluated at 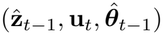. Here (·)^*−*^ denotes the predictive mean and covariance before the measurement correction is applied, and 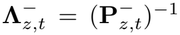 is the corresponding predictive precision.

For the Poisson likelihood, we use a single-step Laplace update at the predicted state mean, with covariance given by the inverse negative Hessian of the log posterior [30], [31]:

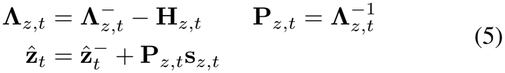

where 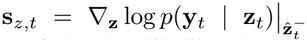 is the state score, the gradient of the log-likelihood at the predicted mean, and 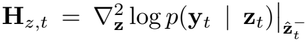 is the corresponding Hessian. For the Poisson log-linear model, the Hessian has the closed form

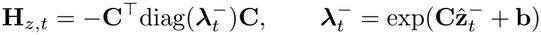

We define the state Fisher information as the Fisher information of the observation likelihood with respect to the latent state **z** :

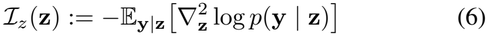

We write 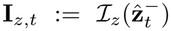, state Fisher information at the predicted state mean at time t. In the Poisson log-linear model, the Hessian is independent of the observed count, so **I**_*z,t*_ = −**H**_*z,t*_. The state score is 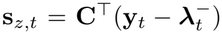. This single-step Laplace update uses the Hessian at the predicted mean rather than at a converged posterior mode; its covarianceform derivation is given in Appendix A.1. An optional iterated refinement recomputes the state score and state Fisher information at each iterate (see (S3) and (S4)).

#### 2) Parameter Update

Individual observations can provide weak parameter information at low signal-to-noise ratio [32], [33]. We therefore accumulate evidence over a block of observations before updating the parameter belief. Within each block, we hold the parameter estimate fixed while updating the latent state at every observation.

Let t_*m*_ and t_*m*+1_ be consecutive update times, *T*_*m*_ = {t_*m*_ + 1, …, t_*m*+1_} the intervening observation block, and n_*m*_ = t_*m*+1_ −t_*m*_ its length. Between updates, the estimate is held fixed, 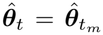 for t_*m*_ ≤t < t_*m*+1_. Each block uses an auxiliary random-walk prior to prevent the approximate posterior from becoming too concentrated while the true parameters remain fixed:

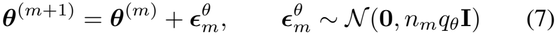

Here ***θ***^(*m*)^ denotes the parameter variable for block m, and q_*θ*_ ≥0 is the covariance increment per observation, so the inflation accumulated over a given number of observations does not depend on the block length. This prior leaves the predictive mean unchanged and inflates the covariance isotropically. At a parameter-update time t, let τ denote the previous update time. Then

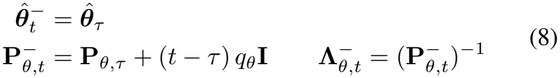

The covariance increment is applied at the end of the block. Let *l*_*t*_(***θ***) = log *p*(**y**_*t*_ | **y**_1:*t−*1_, ***θ***). We use **s**_*θ,t*_ and **H**_*θ,t*_ for local approximations to ∇_***θ***_*l*_*t*_ and 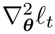, evaluated at the estimate 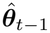 held fixed during the block. For a single increment, the precision-form update is

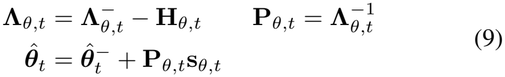

We use Fisher’s and Louis’ identities [34] to express the parameter score and Hessian through the latent-state posterior. We use a Gaussian posterior approximation and neglect parameter derivatives of the predictive covariance 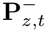. A Gauss–Newton approximation drops second derivatives of the predicted state mean [30], [31]. Let **S**_*t*_ denote the sensitivity of the conditional predicted state mean to ***θ***, with the observation history held fixed. The sensitivity and local increments are

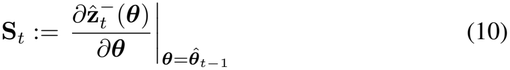

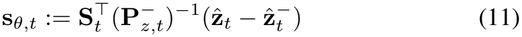

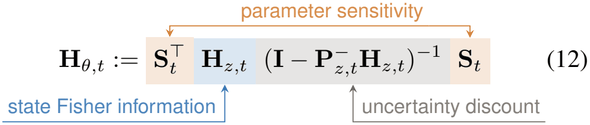

The positive parameter-information approximation is **I**_*θ,t*_ := −**H**_*θ,t*_, which equals 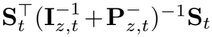 when **I**_*z,t*_ is invertible (S14). A parameter change δ***θ*** shifts the predicted state by **S**_*t*_δ***θ***, and **I**_*θ,t*_ weights this shift using the state Fisher information **I**_*z,t*_ and the predicted state covariance 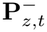. The information is zero when **S**_*t*_δ***θ*** = **0**, for example when δ***θ*** does not change the dynamics along the visited trajectory. In the Poisson log-linear observation model, the information is small where the predicted rates 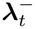 are low or the shift barely changes the log rates **CS**_*t*_δ***θ***. As 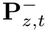 grows, a shift caused by δ***θ*** becomes harder to distinguish from one caused by process noise or state-estimation error, and 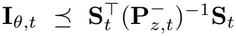. The objective ablations in Section III-D test each factor, and Appendix A.2 gives the one-step derivation.

Because the parameter estimate changes at a block boundary, carrying its sensitivity across that boundary would require replaying the full filtered trajectory under the updated parameter estimate. We treat the filtered state at the start of each block as fixed and reset its parameter sensitivity to zero. We use 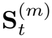 for the block-truncated approximation to the predicted-state sensitivity **S**_*t*_, and 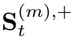 for the corresponding filtered-state sensitivity approximation. With observations held fixed, we propagate

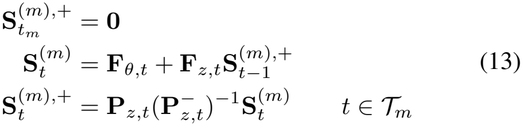

Here **F**_*θ,t*_ = ∂f/∂***θ*** and **F**_*z,t*_ = ∂f/∂**z** are evaluated at 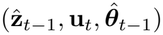. The covariance **P**_*z,t*_ is updated at each observation by (5). We use the predicted sensitivity 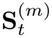 for the current score and Hessian, then carry 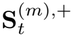 into the next prediction. This block-truncated sensitivity approximation omits parameter derivatives of both predictive and filtered state covariances. Resetting at t_*m*_ also omits parameter dependence inherited from earlier blocks.

We evaluate the local increments in (11) and (12) with 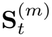 in place of **S**_*t*_ and accumulate them over the block:

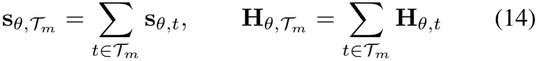

At the end of the block, we apply the predictive prior in (8) with t = t_*m*+1_ and τ = t_*m*_ to update the parameter belief:

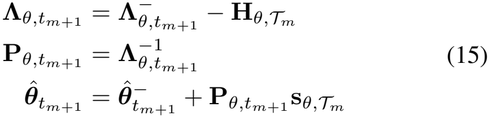

### D. Online Planning

The parameter update in Section II-C2 suggests a direct planning objective: choose inputs that are expected to reduce posterior uncertainty over ***θ***. We therefore formulate input selection as a Bayesian experimental-design problem [25], [26]. At time t, for a candidate input sequence U_*t*+1:*t*+*H*_ := (**u**_*t*+1_, …, **u**_*t*+*H*_), an ideal acquisition objective is the conditional mutual information I(***θ***; **Y**_*t*+1:*t*+*H*_ |*D*_*t*_, U) between the parameters and future observations under that sequence:

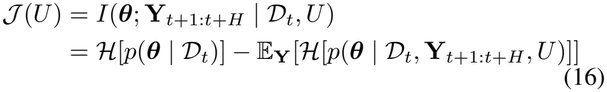

where *D*_*t*_ denotes the data collected up to time t. This expected reduction in parameter posterior entropy is the parameter expected information gain. Maximizing *J*(U) selects the input sequence with the largest expected gain.

The exact objective is intractable as it requires averaging over future observation sequences and updating the parameter posterior for each sequence. We replace this calculation with a tractable local approximation based on deterministic mean rollouts and the parameter-information approximation in (12). We denote this parameter expected information gain (p-EIG) objective by *J*_*θ*_(U).

A tilde denotes prediction under a candidate sequence U, conditioned on the current data *D*_*t*_. During each planning rollout, we hold the parameter mean and covariance fixed and apply no parameter updates or covariance inflation. The rollout starts from the filtered mean:

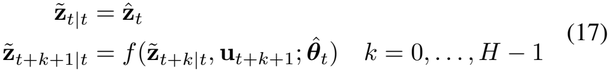

In the following, the rollout index k abbreviates t + k| t. The predicted count mean is 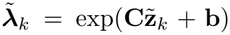. We sample neither process noise nor observations. Inputs are held fixed when differentiating in ***θ***.

For computational simplicity in online planning, we omit measurement corrections to the state covariance and parameter sensitivity within candidate rollouts. Both quantities follow prediction-only recursions; measurement corrections are retained in online inference as actual observations arrive (see (5) and (13)). Initialize 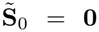 and 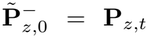. For k = 0, …, H − 1, we propagate

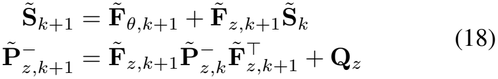

Both Jacobians are evaluated at 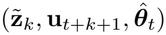.

For k = 1, …, H, the local parameter Hessian approximation is

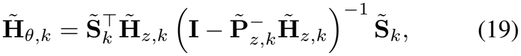

where 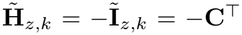 diag 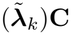 is evaluated at 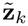, and Ĩ_*z,k*_ is the predicted state Fisher information. Accumulating these local increments gives the approximate terminal precision

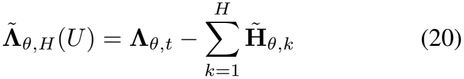

Let 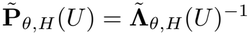. Under the Gaussian parameter belief, posterior entropy equals 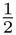 log det **P**_*θ*_ plus a dimension-dependent constant, so the p-EIG objective reduces to a log-determinant reduction in parameter covariance [25], [26]:

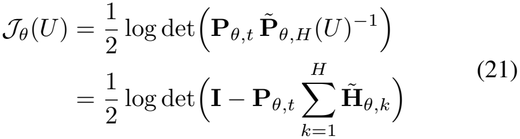

The p-EIG objective is a D-optimal criterion that maximizes parameter information volume [35].

The acquisition objective (21) is non-convex, so we optimize it with iCEM [24], [36],

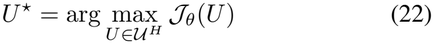

where *U* is the set of admissible inputs. At each decision time, iCEM samples candidate input sequences from a Gaussian distribution, evaluates *J*_*θ*_ using the deterministic rollout above, keeps elite sequences, and updates the proposal mean and scale. After the final iteration, we execute the first K_*u*_ environment-time inputs 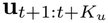 selected by the planner. We update the posterior as observations 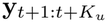 arrive, then replan at t + K_*u*_. This yields a closed-loop procedure that alternates online inference with periodic information-guided input selection. We use this loop as the baseline planner, then add two practical modifications that change the planning time grid and the replanning interval without changing the acquisition objective.

First, coarse-grid planning extends the planning horizon without increasing the number of planning transitions. iCEM searches on a coarser time grid with step cΔt, so that H planning transitions cover cH environment steps; each coarse input is repeated for c environment steps at execution. Second, adaptive intervals allow earlier parameter updates and replanning based on accumulated information or tracking errors. After minimum delays, the parameter update and the replan are triggered early when the posterior-normalized information gain accumulated since the last update exceeds a threshold, or when the filtered state deviates from the planned trajectory by more than a threshold in the metric of the filtering covariance; otherwise they occur at fixed nominal intervals. The rules are given in Appendix A.3.

## III. Experiments and Results

We evaluate PALDI on two neuroscience-motivated multistable systems: a Wilson–Cowan neural mass model of excitatory and inhibitory populations [20] and a decision-making spiking neural network (SNN) [12], [21]. The Wilson–Cowan experiment uses a model-matched simulator, whereas the SNN experiment tests model-class mismatch: agents identify reduced population dynamics from spikes generated by a mechanistic network, using an approximate Poisson observation model. For the SNN, we test both dynamics identification and a decision-control task using the learned model (Section III-E). We ask whether active input selection improves identification, whether the advantage survives degraded observations, and which terms of the acquisition objective account for it.

### A. Evaluation and comparison protocol

To quantify the system recovery, it is tempting to measure the error in parameter recovery, but in nonlinear dynamical systems, the sensitivity of system behavior with respect to parameters is non-trivial at the asymptotic time scale (e.g., small changes can induce a bifurcation). Moreover, the effective behaviors of the system at finite rollout [37], [38] are not generally equally sensitive to each of the governing parameters. We therefore report a global vector-field rollout R^2^, which evaluates how well the learned dynamics reproduce trajectories throughout a prescribed state-space region. Except for the four-region system described below, we sample initial states uniformly in latent space, roll out the true and learned vector fields without input over a fixed horizon, and flatten all latent coordinates into **z**^true^ and 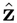. We compute

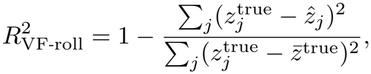

where 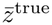 is the mean of the flattened true rollout. This global vector-field rollout score requires access to the true dynamics and is a synthetic-system diagnostic; the spiking-network experiment has no true vector field and uses a held-out rollout score instead (Section III-E). The true and learned rollouts use independent process noise, so the score of the true model itself (the true-model reference in Figure 3A) can fall below one. Negative R^2^ means that a rollout predicts worse than the constant reference 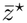.

**Fig. 3.**
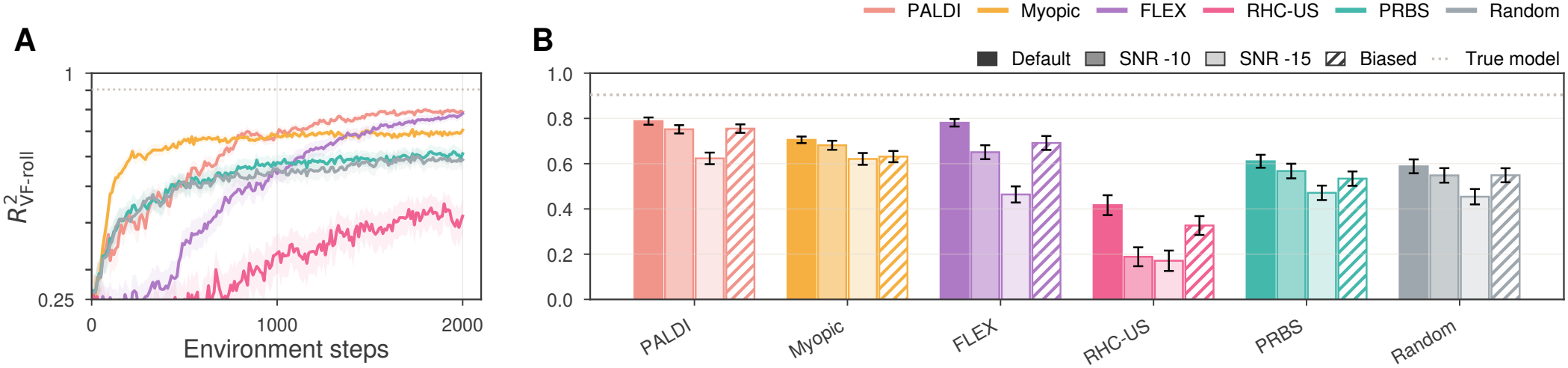
Identification of the neural mass model. Recovery with uniform, weak, and biased observations. (**A**) 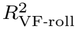 with uniform loading at *−*5 dB; the dotted line marks the true-model reference. (**B**) Final 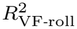 by method under uniform loading at target Fisher SNRs of *−*5, *−*10, and *−*15 dB (dark to light) and under biased loading (hatched), at *−*6 dB. Means and SEM across 100 seeds.

#### Reporting

Within each comparison, all methods share the input budget and evaluation protocol; run settings are given in Appendix B.1. The benchmark figures (Figures 3 and 4) report means and standard errors of the mean (SEM) across runs. Dotted lines show the true-model reference. Because the R^2^ distribution can have a long negative tail, median and interquartile-range (IQR) summaries are provided in Appendix B.8. The spiking-network figure and the appendix diagnostics report medians and 25th–75th percentile IQRs.

**Fig. 4.**
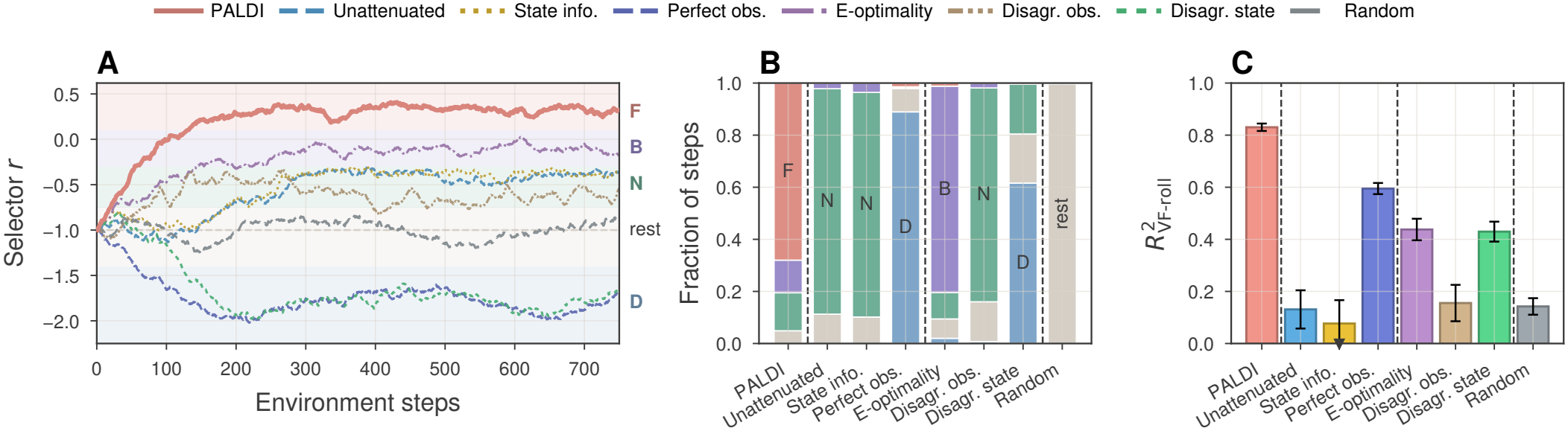
Understanding components of p-EIG. Region preferences and performances across variants (Table I). (**A**) Example selector traces from one matched seed. Shading marks the regions D, N, B, and F, and the dashed line marks the passive equilibrium. (**B**) Mean region occupancy over 750 steps. (**C**) Final rollout prediction performance, averaged over the response coordinates; means and SEM across 100 seeds; error bars that extend below 0 are clipped and marked at the boundary. In (**B**,**C**), dashed lines separate the reference, ablations, variants, and Random.

##### 1) Input-selection baselines

We compare PALDI with baselines that share the same input budget and use the same state-estimation procedure, differing in how they select inputs. Using filtered estimates is necessary because some baselines were originally defined for fully observed systems. Each method otherwise retains its own input-selection rule and, where applicable, its own surrogate-model updates.

#### Non-adaptive inputs

Two conditions select inputs without reference to parameter information. A pseudo-random binary sequence (PRBS) applies inputs at the input bounds and holds each sign pattern for a fixed number of steps, a standard persistently exciting input for system identification [39]. Random open-loop inputs sample each input independently and uniformly within the input bound.

#### Active baselines

Two active baselines optimize an information-related criterion other than the finite-horizon p-EIG: FLEX uses a one-step local D-optimality criterion, whereas RHC-US maximizes finite-horizon predictive variance under a surrogate transition model. Neither criterion accounts for observation information or latent-state uncertainty. FLEX [16] builds its criterion from the local Jacobian of the one-step transition map with respect to the parameter vector under a first-order linearization of the dynamics, 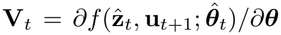. It maintains a Gram matrix **M**_*t*_ of accumulated Jacobians. Its local input-selection rule is based on the D-optimal criterion

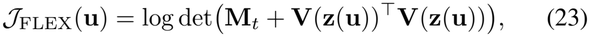

where **z**(**u**) is a locally predicted state. The implementation samples one output coordinate and linearizes its parameter-sensitivity vector in the input, then solves the resulting quadratic program. This approximates the full Gram-matrix criterion. We adapt FLEX by using the EKF-filtered state estimate 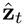 and keeping the noise model fixed. FLEX keeps its own parameter estimator, a regularized least-squares fit to a finite-difference target, in place of the update of Section II-C2. With one-step planning, locally isotropic state Fisher information, and no uncertainty discount, the p-EIG increment has the same sensitivity Gram structure as the FLEX criterion. FLEX thus provides a one-step D-optimal baseline for our finite-horizon p-EIG planner.

The finite-difference target in FLEX amplifies filtering error by 1/Δt, which can destabilize parameter updates. We therefore reject nonfinite or excessively large updates, retaining the previous parameters and Gram matrix. All main comparisons use this stabilized update. Appendix B.3 gives the acceptance limits and compares it with the original update, using the filtered state and the true latent state.

Receding Horizon Curiosity with uncertainty sampling (RHC-US) [40] selects inputs by maximizing predictive uncertainty under a surrogate transition model rather than parameter information. It models state increments with a Bayesian linear model on feature embeddings, Δ**z**_*t*+1_ := **z**_*t*+1_ −**z**_*t*_ = **W**^⊤^ψ([**z**_*t*_, **u**_*t*+1_]), and maintains a posterior over **W**. The input sequence is chosen over a finite horizon by maximizing predictive variance,

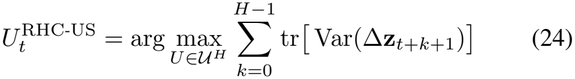

In the original method, the selected sequence is executed openloop and the surrogate model is updated only after rollout. For our setting, we use the EKF-filtered latent-state estimate instead of the true state and treat one planning horizon as one episode: the planned sequence is executed open-loop, and the surrogate model is updated only after the horizon. The surrogate only selects inputs; the evaluated model is the parameter estimate of Section II-C2.

#### Myopic

Myopic evaluates observation information at one-step future state, whereas the main PALDI plans over a longer horizon with adaptive update and replanning intervals (Appendix B.1). The objective ablations in Section III-D vary only the acquisition objective (Table I).

**TABLE 1.** Acquisition objectives up to scaling: reference, ablations, and variants. All objectives are maximized over *U*. eig_min_ denotes the smallest eigenvalue. 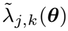 and 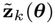 denote predicted mean observation and states

| **Objective** | **Acquisition objective** |
|---|---|
| PALDI p-EIG | $\log\det\left(\mathbf{I} - \mathbf{P}_{\theta,t}\sum_{k=1}^{H}\tilde{\mathbf{H}}_{\theta,k}\right)$ |
| Unattenuated p-EIG<br>(−state uncertainty) | $\log\det\left(\mathbf{I} - \mathbf{P}_{\theta,t}\sum_{k=1}^{H}\tilde{\mathbf{S}}_k^\top\tilde{\mathbf{H}}_{z,k}\tilde{\mathbf{S}}_k\right)$ |
| Perfect-obs. p-EIG<br>(−state Fisher) | $\log\det\left(\mathbf{I} + \mathbf{P}_{\theta,t}\sum_{k=1}^{H}\tilde{\mathbf{S}}_k^\top(\tilde{\mathbf{P}}_{z,k}^{-})^{-1}\tilde{\mathbf{S}}_k\right)$ |
| State Information<br>(−param. sensitivity) | $\sum_{k=1}^{H}\log\det\left(\mathbf{I} - \tilde{\mathbf{P}}_{z,k}^{-}\tilde{\mathbf{H}}_{z,k}\right)$ |
| E-optimality | $\mathrm{eig}_{\min}\left(-\mathbf{P}_{\theta,t}\sum_{k=1}^{H}\tilde{\mathbf{H}}_{\theta,k}\right)$ |
| Disagreement in obs. | $\sum_{k=1}^{H}\sum_{j}\log\left(1 + \mathrm{Var}_{\boldsymbol{\theta}\sim q_t}[\tilde{\lambda}_{j,k}(\boldsymbol{\theta})]\right)$ |
| Disagreement in state | $\sum_{k=1}^{H}\mathrm{tr}\,\mathrm{Cov}_{\boldsymbol{\theta}\sim q_t}[\tilde{\mathbf{z}}_k(\boldsymbol{\theta})]$ |

### B. System identification with information-guided inputs

A planner can collect useful data by moving the system beyond the region it explores without input. On a bistable Wilson–Cowan population model, PALDI recovers the dynamics more accurately than the other policies, and it explores more of the state space (Figures S1 and 3). Without input, trajectories settle near one of two stable states, and the resulting observations provide little information about the parameters governing dynamics in unvisited regions.

The Wilson–Cowan system [20] is a neural mass model of an excitatory and an inhibitory population with firing rates E_*t*_ and I_*t*_ (Figure 2A). With refractory factors and time in units of the membrane time constant, the drift components are

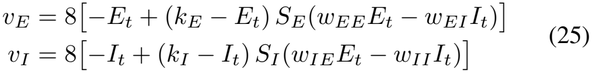

where **v**(**z**_*t*_; ***θ***) = (v_*E*_, v_*I*_)^⊤^ is the drift, and S_*E*_ and S_*I*_ are sigmoid gain functions with S_*j*_(0) = 0. The parameters ***θ*** are the four synaptic weights. We use a standard parameter set with two stable states [41]; its values are given in Appendix B.1. The circuit has a stable down state at E = I = 0 and a stable up state, and the stable manifold of a saddle separates their basins (Figure 2B). Without noise or external input, a trajectory remains in its initial basin. The inputs u_*E,t*_ and u_*I,t*_ act on the excitatory and inhibitory populations, a simplified model of stimulation that targets each cell type, and can drive the state across the basin boundary (Figure 2B,F). The latent state **z**_*t*_, a centered and rescaled (E_*t*_, I_*t*_), evolves as

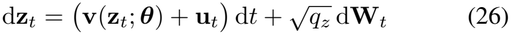

where **W**_*t*_ is a standard Wiener process. The discretization is given in Appendix B.1.

Figure 2C shows a map of ∥∂**v**/∂θ_*i*_∥ for each parameter on a shared color scale; larger values indicate a greater change in drift for a small change in that parameter. At the down state, E_*t*_ = I_*t*_ = 0 and the parameter sensitivity is zero. Activity near this state is therefore weakly informative about the parameters, motivating inputs that move the system away from it. The parameters are positive by definition, and every method starts from initial parameter estimates with the correct signs (Appendix B.1).

We observe the state through d_*y*_ = 20 Poisson neurons with rates **r**_*t*_ = exp(**Cz**_*t*_ + **b**_*r*_) in Hz. The loading directions are spread evenly around the latent plane, and NeuroFisherSNR [42] calibrates the bias and loading to a target Fisher SNR of −5 dB with mean and maximum rates of 20 and 100 Hz (Appendix B.1). Figure 2D–F shows the loading directions, the resulting state Fisher information, and example spike trains for the observation models of Section III-C.

On this system, PALDI reaches a mean 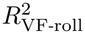 of 0.79; the true-model reference is about 0.9 (Figure 3A). FLEX reaches 0.78, Myopic 0.71, PRBS 0.61, random inputs 0.59, and RHC-US 0.42. PALDI and FLEX do not differ significantly, but FLEX improves more slowly and reaches the accuracy of PALDI only near the end of the run. The lower plateaus of Myopic and non-adaptive inputs are consistent with limited exploration beyond the local attractor and lower parameter information gain (Appendix B.2). These results suggest that PALDI’s information-guided planning is useful when identification requires reaching informative regions beyond those visited without input.

For FLEX, both the state estimate and the update rule affect this comparison. Here it uses the EKF-filtered state 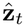 and the stabilized parameter update described in Section III-A1.

With the filtered state and its original update, FLEX fails in every run (mean −1.85, SEM 0.07). With the true latent state and the original update, the mean rises only to −0.04 (0.11) (Appendix B.3). Access to the true state improves FLEX’s median performance, but the performance remains less robust across seeds.

### C. Robustness to observation degradation

We next test the methods under two realistic recording conditions in which observations of the bistable model are weak or carry little information about one population. First, we lower the target Fisher SNR from −5 to −10 and −15 dB, which reduces the state Fisher information throughout. Second, we bias the loading directions toward the inhibitory axis, so that the observations carry little information about the excitatory population (biased loading; Figure 2D,E). All but one of the 20 loading directions lie within about 50^*°*^ of the inhibitory axis (Appendix B.1).

Again, PALDI has the highest mean final 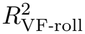 among the main comparison policies in every condition (Figure 3B). FLEX ties PALDI at high SNR but degrades more quickly than PALDI as the SNR decreases. Myopic declines less with SNR, but it is overall less accurate than PALDI, and the gap is largest under biased loading.

Myopic and PALDI both use observation information to select inputs, Myopic over one step and PALDI over a finite horizon. Planning over the horizon adds accuracy where observations are uniform and moderately noisy and, most of all, under biased loading, where the observations inform one population poorly. At −15 dB the two methods tie, so the longer horizon gives no measurable advantage at the lowest SNR. The objectives of FLEX and RHC-US do not account for observation information. FLEX matches PALDI when the observations are strongest; FLEX and RHC-US both lose more accuracy than the other methods as the SNR decreases. These results suggest that an identification objective should account for the information in the observations, and that planning ahead with such an objective adds accuracy except at the lowest SNR.

### D. Effect of the acquisition objective on exploration

The proposed p-EIG objective combines the three components of (12): (A) parameter sensitivity, the sensitivity of the state trajectory to the dynamics parameters; (B) state Fisher information, the sensitivity of observations to latent states; and (C) the uncertainty discount due to latent-state uncertainty. Without any one of them (ablations in Table I), the planner may fail to correctly attribute changes in the observations to changes in the parameters and explore less informative regions. To understand and demonstrate the role of each component, we designed an eight-dimensional four-region system, **z** = (r, **s, b**, h), with two responses **s** = (s_1_, s_2_, s_3_) and **b** = (b_1_, b_2_, b_3_). Along the first state dimension *r* (the selector), inputs move the state from rest into four regions (D, N, B, and F) of distinct dynamics; region D lies on the opposite side of rest from the other three. The latent state is governed by

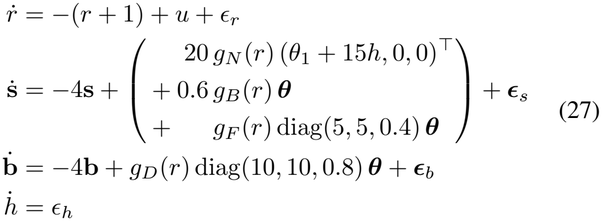

where ϵ_*r*_, ***ϵ***_*s*_, ***ϵ***_*b*_, and ϵ_*h*_ are white process noise. Note that the parameters ***θ*** only affect the response dynamics 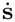 and 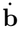, and the input u only drives the selector r. Along r, the radial basis functions g_*D*_(r), g_*N*_ (r), g_*B*_(r), and g_*F*_ (r) define regions with different parameter sensitivity. All coordinates are observed through noisy Poisson spike counts, but the observations are much less sensitive to the state h, so its estimate stays uncertain, and to the response **b**, so a change in **b** is hard to detect. In region N (nuisance), θ_1_ and h drive s_1_, both with a strong gain. In regions B (balanced) and F (focused), all three parameters drive **s**: weakly but equally in B, and strongly through θ_1_ and θ_2_ but weakly through θ_3_ in F. In region D (dim), all three parameters drive the second response **b** with twice the gain of region F, but the observations are less sensitive to **b** than to **s**. Details of the regions, the observation model, and the experiment are given in Appendix B.4.

In Figure 4, all methods start at rest, but only PALDI mostly samples region F (panels A and B) and achieves the highest mean 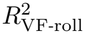 (panel C). The ablations instead sample in less informative regions. Unattenuated p-EIG and State Information sample predominantly in region N, misled by the high variance of s_1_ due to the high gain and the uncertain h, although this region carries no information about θ_2_ and θ_3_. Perfect-observation p-EIG keeps the uncertainty discount and avoids region N, but it stays in region D, where the parameters act strongly on coordinates that the observations barely explain.

Alongside the component ablations, we compare alternative objectives (Table I). E-optimality uses the same parameter information as PALDI and therefore successfully leaves region N. However, since it maximizes the information in the least-informed direction, it avoids region F, where θ_3_ gains little information, and stays mostly in region B (Figure 4B). Disagreement-based objectives that maximize the variance of predicted rates or states with respect to the parameter uncertainty (Appendix B.4) serve as a proxy for information gain [43] but do not account for uncertain latent states. As in the noisy-TV problem [44], the observation-based variant favors the high-variance response in region N despite its limited parameter information. The state-based variant mostly favors region D, where the predicted response is largest.

These results demonstrate the importance of combining parameter sensitivity, state Fisher information, and the uncertainty discount in the acquisition objective: objectives that omit any one of them, or that weigh only the least-informed parameter direction, can favor less informative regions and recover the dynamics less accurately.

### E. Model learning for efficient control under model mismatch

The preceding benchmarks show how active input selection affects identification when the agent uses the same dynamics and observation model classes as the data-generating system. To test whether these benefits extend to a more realistic neural system, we use a decision-making circuit implemented as a spiking neural network, where both the reduced dynamics and the Poisson observation model are approximations. The question is whether active identification can still learn a useful predictive model, and whether that model is accurate enough to improve control of the circuit’s decisions.

#### Circuit and reduced model

We simulate a decision network [21] in Brian 2 [45]. Its two selective excitatory pools compete through shared inhibition, and the inputs inject current into each pool (Figure S3).

As the parametric dynamical description, we use a reduced model [12], whose two states are the slow, population-averaged NMDA gating variables s_1_ and s_2_, with faster network variables approximated by their equilibrium responses. With centered and rescaled states 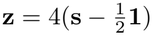, the reduced drift is

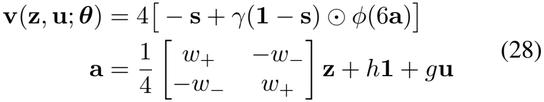

Here ϕ(q) = q/(1− e^*−q*^), with ϕ(0) = 1, acts elementwise⊙, denotes elementwise multiplication, and **1** = (1, 1)^⊤^. The first term describes NMDA decay; the second describes activation, limited by the fraction 1 −s_*i*_ of closed gates. The vector **a** is the effective drive to the two pools, combining recurrent activity, a constant background drive, and the applied input. Each agent estimates the five coefficients shared by the pools: self-excitation w_+_, cross-inhibition w_*−*_, background drive h, activation scale γ > 0, and input gain g > 0. These coefficients are learned as effective parameters of the reduced model. The parameterization and the separately fitted reference model are given in Appendix B.6.

We observe the circuit through spike counts from 20 of the 240 excitatory neurons in each pool, giving 40 recorded neurons in total, in 5-ms bins. To link these counts to the reduced state, we fit the Poisson readout **C, b**_*r*_ (2) using paired population states and spike counts from separate simulations.

The readout then remains fixed while each agent estimates states and dynamics parameters from spikes, without access to the recorded population states (Appendix B.5).

#### Identification

Each identification trial starts from the reset state **z** = (− 2, −2)^⊤^, and the agent applies inputs while observing spikes. A decision occurs when one pool leads the other by |z_1_ −z_2_| > 2. The first five trials simulate a classic decision-making experiment: the circuit receives sensory evidence, 6 pA to a randomly selected pool during the first second, and it resets 200 ms after the decision or at the 2-s trial limit. In these trials, the agent’s input is limited to 2 pA, below the evidence, so the decision follows the evidence. The last fifteen trials have no evidence, allow inputs up to 10 pA, and always last 2 s, so the agent keeps driving the circuit after its decision. Agents know the evidence, learn of each reset when it occurs, and retain their learned dynamics across trials. We evaluate how accurately each learned model predicts population activity over 500 ms in network simulations not used for identification, measured by 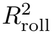. Each prediction starts from the recorded population state and receives no subsequent corrections from spikes. For comparison, we fit the same reduced model directly to recorded population states from separate simulations. Details are given in Appendices B.5 and B.6.

PALDI learns the most accurate model among the identification policies. Its median held-out 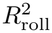 rises from 0.70 after the five evidence trials to 0.91 after twenty, close to the separately fitted reference of 0.97 (Figure 5C). Myopic, RHC-US, and FLEX reach 0.86, 0.77, and 0.58, respectively. PRBS and random inputs reach 0.58 and 0.55 after the evidence trials but fall to 0.19 and 0.11 after twenty—the additional trials do not improve every identification method. Of the six methods, only PALDI and Myopic outperform the passive learner, which receives the evidence in all twenty trials and applies no input.

**Fig. 5.**
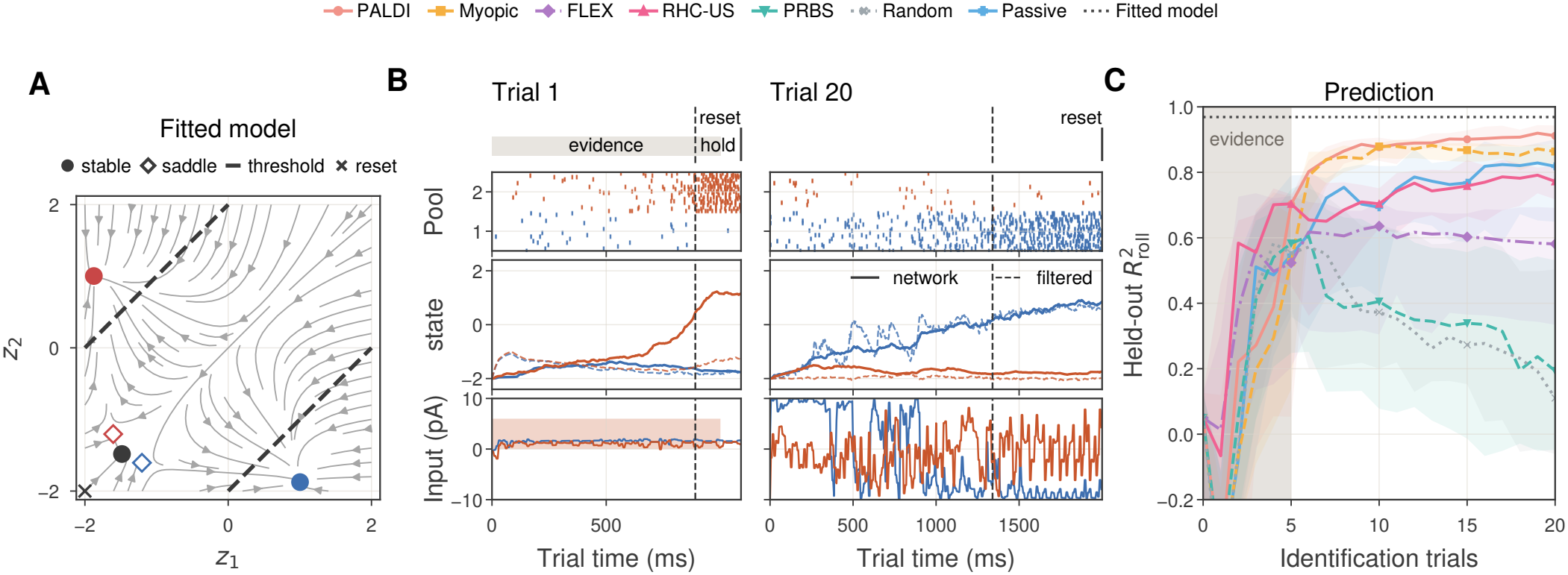
Identification of the spiking network under model mismatch. Reduced-model dynamics, example identification trials, and prediction of network activity. (**A**) Input-free phase portrait of the reduced model fitted to recorded population states. Markers show stable fixed points (circles) and saddles (diamonds), colored by pool (blue, 1; red, 2; black, undecided), and the reset state (cross); dashed lines mark the decision thresholds |*z*_1_ *−z*_2_| = 2. (**B**) Trials 1 and 20 of PALDI: spike counts, network states (solid), filtered states (dashed), and inputs (pA per neuron). The gray bar marks 6 pA evidence in trial 1, and dashed lines decisions; in trial 1 the agent’s input is limited to 2 pA. (**C**) Rollout prediction performance over the identification trials; medians and IQR across 20 training seeds; the band marks the five evidence trials.

#### Control

We next test whether the learned dynamics can support effective control of the system. The task is to induce a change of mind. We call the pool that wins the initial decision the *chosen pool* (c) and the other pool the *alternative pool* (a). To make the alternative pool win, the controller must drive the network out of the attractor of the chosen pool and across the saddle (Figure 5A); doing this efficiently requires an accurate model of the whole landscape, including both attractors, the saddle, and the flow between them. The network receives an external evidence current in one randomly selected pool during the first second and makes its initial decision without control. Then, the controller applies pers, with ∥**i**_*t*_∥_2_ ≤ 20 pA and a total budget 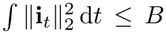, neuron currents **i**_*t*_ = (i_1,*t*_, i_2,*t*_)^*⊤*^ to the two pools for up to 2 where B = 480 pA^2^ s (other budgets in Figure S4). We freeze each agent’s learned parameters and use its model in the same iCEM controller. Success means that the network reverses its decision, z_*a*_ −z_*c*_ > 2, within 2.5 s of the initial decision (Figure 6A). We compare models learned after different numbers of identification trials with the fitted reference model and fixed open-loop inputs (Appendix B.5).

**Fig. 6.**
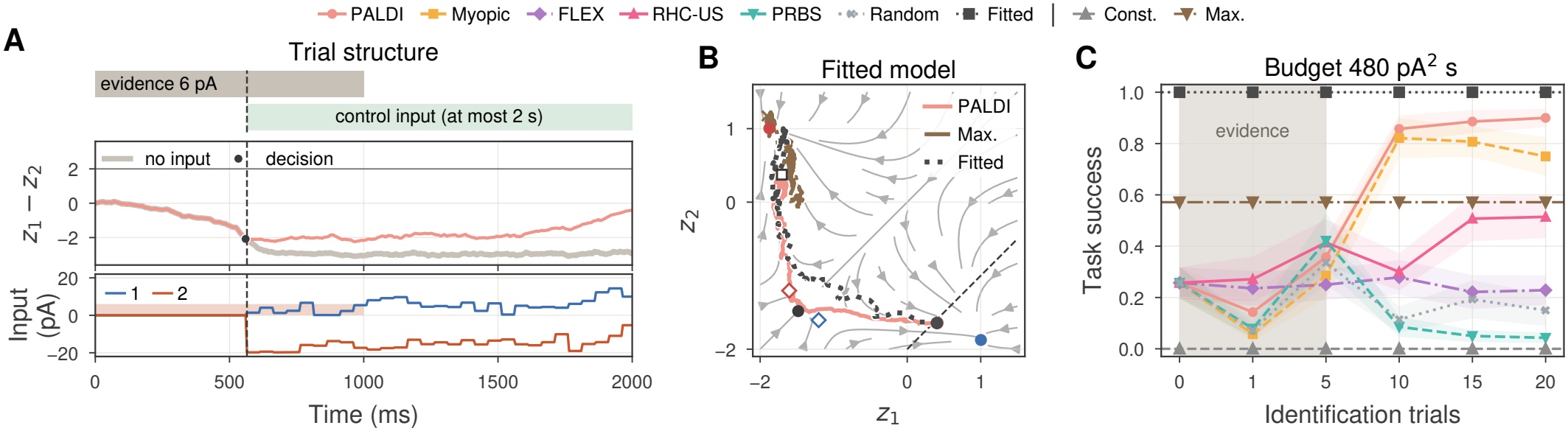
Control of the spiking network with learned models. Task, example trajectories, and success across trials. (**A**) Evidence (gray bar, 6 pA) leads to an initial decision (circle, vertical dashed line); the controller then has at most 2 s (green) to drive *z*_1_*− z*_2_ across the reversal threshold (horizontal line) with currents to pools 1 and 2. The gray trace shows the trial without control. (**B**) Controlled trajectories in the trial of (**A**) with PALDI, Maximum-amplitude, and the fitted model (480 pA^2^ s), over the phase portrait of Figure 5A. Markers show control onset (square), successful termination (circle), and the unsuccessful endpoint (cross); the dashed line is the reversal threshold *z*_1_ *−z*_2_ = 2 (initial choice: pool 2). (**C**) Success across identification trials at 480 pA^2^ s with the models learned by the different methods; the band marks the evidence trials. Const. and Max. denote the fixed open-loop Constant-and Maximum-amplitude inputs.

In addition to the previous baselines, we compare two fixed open-loop inputs: Maximum-amplitude applies the maximum input from control onset until the budget is spent, whereas Constant-amplitude spreads the budget evenly over the control window. Both excite the alternative pool and suppress the chosen pool. In the example trial in Figure 6B, the controller with the PALDI model reaches the reversal threshold, while Maximum-amplitude does not.

After twenty trials, the PALDI model achieves the highest success rate among the models learned by the different agents (Figure 6C). With the same control budget, the success rate depends on how the current is delivered. Spreading it evenly (Constant-amplitude) never reverses the decision, and pushing as hard as possible (Maximum-amplitude) gives a success rate of 0.57. The iCEM controller with the PALDI model, adaptively replanning and shaping the currents, reaches 0.90 after twenty identification trials, compared with 1.00 for the fitted reference model; the next-best learned model (Myopic) reaches 0.75. However, PALDI requires a larger control budget to approach the fitted model’s success rate, consistent with residual model error (Figure S4). These results show that learned dynamics support effective control in the change-of-mind task despite model-class mismatch.

## IV. Discussion

We proposed PALDI, a novel information-guided finitehorizon planning method that improves online identification of nonlinear latent dynamics under partial observability. It addresses a common problem in neuroscience where a system of interest rests in one of the stable states with only local fluctuations [46].

In the neural mass model experiment, PALDI identified the dynamics nearly as accurately as the true bistable model (Section III-B). PALDI also addresses another common problem in neuroscience: experimentalists often have access to an idiosyncratic subset of neurons that may have different firing rate, tuning curve, and noise. PALDI stayed accurate when observations had a low signal-to-noise ratio or were less informative in some regions of the state space (Section III-C). We showed the effectiveness of information-guided finite-horizon planning for identifying latent dynamics when attractors confine activity and the information about the parameters is unevenly distributed across the state space—PALDI plans inputs that steer the system out of the attractor and into the regions whose observations reveal the most about the parameters.

The four-region system illustrates the role of each component of the p-EIG objective (Section III-D). Objectives that omit parameter sensitivity or the uncertainty discount favored a region where the observation varies mainly because of uncertainty in the state rather than the parameters. An objective that omits the state Fisher information favored a region where the parameters act strongly on states that the observations barely explain. All of them recovered the dynamics less accurately than PALDI. Input design under partial observability should therefore account for how the parameters can alter the state trajectory, how well the observations resolve the state, and how uncertain the state estimate is.

In the spiking decision network, both the reduced dynamics and the Poisson readout only approximate the data-generating circuit (Section III-E). PALDI still learned an accurate model with high prediction accuracy. In the change-of-mind task, a model-based controller used each learned model to plan inputs that reverse the circuit’s initial decision. Under the same control budget, the model learned by PALDI reversed the decision more reliably than the models learned by the other identification policies. Active identification can thus yield a model that is useful for control despite model mismatch. This suggests a potential contribution of PALDI to model-based closed-loop stimulation [6], [8], from experimental design to adaptive therapies such as adaptive deep brain stimulation and responsive neurostimulation, which currently adjust stimulation with threshold or detection rules rather than a model [4], [5].

### Relation to prior work

These findings agree with prior work showing that active input design can improve identification over passive data collection [16]–[19], and extend this pattern to partially observed systems that may need inputs planned over several steps to leave an attractor. p-EIG is a tractable online approximation to sequential expected information gain [25]–[27], and its finite-horizon planning connects to planning-to-explore methods [43], [47]–[49]. Concurrently, Arbelaiz and colleagues [50] developed a sampling-based, information-guided planning algorithm for fully observed systems.

### Limitations and future directions

We made a number of explicit assumptions and approximations in favor of tractability and computational efficiency. Gaussian state and parameter beliefs and EKF cannot represent multimodal posteriors nor accommodate strongly nonlinear dynamics [1]–[3]. The parameter update omits covariance derivatives and second-order dynamics terms, and planning uses prediction-only covariance propagation on a coarse time grid; the error of p-EIG relative to mutual information remains to be quantified. The observation model is assumed known; learning it jointly with the dynamics from spikes is a natural extension. Likewise, the functional form of the latent dynamics is assumed known. Expressive models, such as neural networks, can represent more flexible dynamics, but their high-dimensional parameter space makes p-EIG, whose cost grows at least quadratically with the parameter dimension, impractical for real-time planning. A possible future direction is to combine PALDI with models that represent related systems with shared dynamics and a low-dimensional, system-specific embedding [51], so that only the low-dimensional embedding needs to be identified online. Closed-loop experiments will also require planning within the latency of the recording and stimulation hardware.

