## Supplementary Material for "Information-Guided Planning for Online System Identification of Partially Observed Nonlinear Dynamical Systems"

This Supplementary Material contains Appendices A and B of the main text: the derivations, additional experimental results, and experimental settings referenced there. Equations, figures, and tables numbered with the prefix S are defined here; numbers without the prefix, and section numbers such as Section III-B, refer to the main text.

#### Appendix A
#### Inference and planning details

##### A.1 State update

The single-step Laplace update in (5) has an equivalent covariance form when the residual is expressed in natural-parameter coordinates. Let $h$ map the latent state to the natural parameter $\boldsymbol{\eta} = h(\mathbf{z})$ of the observation distribution ($h(\mathbf{z}) = \mathbf{C}\mathbf{z} + \mathbf{b}$ in the Poisson log-linear model), and let $\boldsymbol{\eta}_t^- = h(\hat{\mathbf{z}}_t^-)$, $\mathbf{s}_{\eta,t} = \nabla_{\boldsymbol{\eta}} \log p(\mathbf{y}_t \mid \boldsymbol{\eta})\big|_{\boldsymbol{\eta}_t^-}$, and $\mathbf{I}_{\eta,t} = \mathcal{I}_\eta(\boldsymbol{\eta}_t^-)$. Write

$$\mathbf{D}_t = \frac{\partial h}{\partial \mathbf{z}}\Big|_{\hat{\mathbf{z}}_t^-}, \qquad \boldsymbol{\nu}_t = \mathbf{I}_{\eta,t}^{-1}\mathbf{s}_{\eta,t}$$

Here $\boldsymbol{\nu}_t$ is a pseudo-observation residual, not a raw count residual. For the Poisson log-linear model,

$$\mathbf{D}_t = \mathbf{C}, \quad \mathbf{I}_{\eta,t} = \operatorname{diag}(\boldsymbol{\lambda}_t^-), \quad \boldsymbol{\nu}_t = \operatorname{diag}(\boldsymbol{\lambda}_t^-)^{-1}(\mathbf{y}_t - \boldsymbol{\lambda}_t^-)$$

Using $\mathbf{P}_{\nu,t}$ for the innovation covariance of this local Gaussian pseudo-observation model gives

$$\begin{aligned}
\mathbf{P}_{\nu,t} &= \mathbf{D}_t\mathbf{P}_{z,t}^-\mathbf{D}_t^\top + \mathbf{I}_{\eta,t}^{-1} \\
\mathbf{K}_t &= \mathbf{P}_{z,t}^-\mathbf{D}_t^\top\mathbf{P}_{\nu,t}^{-1} \\
\hat{\mathbf{z}}_t &= \hat{\mathbf{z}}_t^- + \mathbf{K}_t\boldsymbol{\nu}_t \\
\mathbf{P}_{z,t} &= (\mathbf{I} - \mathbf{K}_t\mathbf{D}_t)\mathbf{P}_{z,t}^-
\end{aligned} \tag{S1}$$

In particular, $\mathbf{K}_t\boldsymbol{\nu}_t = \mathbf{P}_{z,t}\mathbf{C}^\top(\mathbf{y}_t - \boldsymbol{\lambda}_t^-)$, which recovers the state mean update in (5). The mean correction vanishes when the observed counts equal their predicted means.

**State Fisher information and observed information.** For a differentiable observation map, the state Fisher information is given exactly by

$$\mathbf{I}_{z,t} := \mathcal{I}_z(\hat{\mathbf{z}}_t^-) = \mathbf{D}_t^\top\mathbf{I}_{\eta,t}\mathbf{D}_t \tag{S2}$$

For nonlinear $h$, the observed information about the state also contains terms involving second derivatives of $h$, weighted by the natural-parameter score. It can be indefinite. Replacing this observed information with state Fisher information is an approximation. For the Poisson log-linear model, $h$ is affine and the natural-parameter Hessian is independent of the observed count, so $\mathbf{I}_{z,t} = -\mathbf{H}_{z,t} = \mathbf{C}^\top \operatorname{diag}(\boldsymbol{\lambda}_t^-)\mathbf{C}$ exactly at the evaluation point.

**Optional iterated refinement.** We use the single-step Laplace update in (5). A refinement can instead start at $\hat{\mathbf{z}}_t^{(0)} = \hat{\mathbf{z}}_t^-$ and recompute the state score $\mathbf{s}_{z,t}^{(i)}$ and state Fisher information $\mathbf{I}_{z,t}^{(i)} = \mathcal{I}_z(\hat{\mathbf{z}}_t^{(i)})$ at each iterate. The Fisher-scoring step solves

$$\left(\boldsymbol{\Lambda}_{z,t}^- + \mathbf{I}_{z,t}^{(i)}\right)\Delta\mathbf{z}_t^{(i)} = \mathbf{s}_{z,t}^{(i)} - \boldsymbol{\Lambda}_{z,t}^-(\hat{\mathbf{z}}_t^{(i)} - \hat{\mathbf{z}}_t^-), \tag{S3}$$

then sets $\hat{\mathbf{z}}_t^{(i+1)} = \hat{\mathbf{z}}_t^{(i)} + \Delta\mathbf{z}_t^{(i)}$. For the Poisson log-linear model, this is a Newton step because state Fisher information and observed information coincide. After $k$ iterations, recompute the state Fisher information at the final iterate and set

$$\hat{\mathbf{z}}_t = \hat{\mathbf{z}}_t^{(k)}, \qquad \boldsymbol{\Lambda}_{z,t} = \boldsymbol{\Lambda}_{z,t}^- + \mathbf{I}_{z,t}^{(k)} \tag{S4}$$

Only after convergence does this give the usual Laplace approximation at a posterior mode.

##### *A.2 Parameter update*

We derive the local parameter score and Hessian in (11) and (12) for a single transition. We start from the exact marginal log-likelihood

$$\ell_t(\boldsymbol{\theta}) = \log p(\mathbf{y}_t \mid \mathbf{y}_{1:t-1}, \boldsymbol{\theta}) = \log \int p(\mathbf{y}_t \mid \mathbf{z}_t)\, p(\mathbf{z}_t \mid \mathbf{y}_{1:t-1}, \boldsymbol{\theta})\, d\mathbf{z}_t \tag{S5}$$

Let $p_t(\mathbf{z}_t) = p(\mathbf{z}_t \mid \mathbf{y}_{1:t}, \boldsymbol{\theta})$ denote the filtering posterior. Since the Poisson observation likelihood does not depend on $\boldsymbol{\theta}$ directly, Fisher's identity [1] gives

$$\nabla_{\boldsymbol{\theta}} \ell_t(\boldsymbol{\theta}) = \mathbb{E}_{p_t}[\nabla_{\boldsymbol{\theta}} \log p(\mathbf{z}_t \mid \mathbf{y}_{1:t-1}, \boldsymbol{\theta})]\,, \tag{S6}$$

and Louis' identity [1] gives the observed information (complete minus missing)

$$-\nabla^2_{\boldsymbol{\theta}} \ell_t(\boldsymbol{\theta}) = \mathbb{E}_{p_t}\left[-\nabla^2_{\boldsymbol{\theta}} \log p(\mathbf{z}_t \mid \mathbf{y}_{1:t-1}, \boldsymbol{\theta})\right] - \mathrm{Cov}_{p_t}(\nabla_{\boldsymbol{\theta}} \log p(\mathbf{z}_t \mid \mathbf{y}_{1:t-1}, \boldsymbol{\theta})) \tag{S7}$$

**Gaussian state prediction.** We use the local Gaussian model

$$p(\mathbf{z}_t \mid \mathbf{y}_{1:t-1}, \boldsymbol{\theta}) = \mathcal{N}(\mathbf{z}_t; \hat{\mathbf{z}}_t^-(\boldsymbol{\theta}), \mathbf{P}_{z,t}^-), \qquad \hat{\mathbf{z}}_t^-(\boldsymbol{\theta}) = f(\hat{\mathbf{z}}_{t-1}, \mathbf{u}_t; \boldsymbol{\theta}), \tag{S8}$$

with $\mathbf{P}_{z,t}^-$ and $\hat{\mathbf{z}}_{t-1}$ held fixed when differentiating in $\boldsymbol{\theta}$. We evaluate the derivatives at $\hat{\boldsymbol{\theta}}_{t-1}$, the parameter estimate held fixed during the block, which equals the prior mean $\hat{\boldsymbol{\theta}}_t^-$ of (8), and use the sensitivity $\mathbf{S}_t$ defined in (10). Under these one-step assumptions, $\mathbf{S}_t = \mathbf{F}_{\theta,t}$. Then

$$\nabla_{\boldsymbol{\theta}} \log p(\mathbf{z}_t \mid \cdot, \boldsymbol{\theta})\big|_{\hat{\boldsymbol{\theta}}_{t-1}} = \mathbf{S}_t^\top (\mathbf{P}_{z,t}^-)^{-1} (\mathbf{z}_t - \hat{\mathbf{z}}_t^-) \tag{S9}$$

$$-\nabla^2_{\boldsymbol{\theta}} \log p(\mathbf{z}_t \mid \cdot, \boldsymbol{\theta})\big|_{\hat{\boldsymbol{\theta}}_{t-1}} = \mathbf{S}_t^\top (\mathbf{P}_{z,t}^-)^{-1} \mathbf{S}_t \;-\; \sum_{i=1}^{d_z} \left[(\mathbf{P}_{z,t}^-)^{-1}(\mathbf{z}_t - \hat{\mathbf{z}}_t^-)\right]_i \, \nabla^2_{\boldsymbol{\theta}} \hat{z}_{t,i}^-(\boldsymbol{\theta})\big|_{\hat{\boldsymbol{\theta}}_{t-1}} \tag{S10}$$

**Single-step Laplace update.** Both identities take expectations under the filtering posterior $p_t$, which is unavailable in closed form. The state update of Appendix A.1 supplies a Gaussian posterior approximation,

$$p_t(\mathbf{z}_t) \approx \mathcal{N}(\mathbf{z}_t; \hat{\mathbf{z}}_t, \mathbf{P}_{z,t}), \qquad \mathbf{P}_{z,t} = \left((\mathbf{P}_{z,t}^-)^{-1} + \mathbf{I}_{z,t}\right)^{-1}, \tag{S11}$$

whose mean is the single corrected state estimate and whose covariance uses the Hessian at the predicted state mean, as in (5). If the optional iterations converge, both can instead be evaluated at the posterior mode. Substituting its first two moments into the score and information identities, and evaluating at $\boldsymbol{\theta} = \hat{\boldsymbol{\theta}}_{t-1}$, produces the quantities used in (9).

**Local parameter score.** Under the fixed-covariance Gaussian prior, Fisher's identity needs only the posterior mean. Approximating $\mathbb{E}_{p_t}[\mathbf{z}_t] \approx \hat{\mathbf{z}}_t$ gives

$$\mathbf{s}_{\theta,t} := \mathbf{S}_t^\top (\mathbf{P}_{z,t}^-)^{-1} (\hat{\mathbf{z}}_t - \hat{\mathbf{z}}_t^-), \qquad \hat{\mathbf{z}}_t^- = \hat{\mathbf{z}}_t^-(\hat{\boldsymbol{\theta}}_{t-1}), \tag{S12}$$

which is (11): the state correction $\hat{\mathbf{z}}_t - \hat{\mathbf{z}}_t^-$ due to the observation, weighted by the predictive state precision and mapped into parameter space by $\mathbf{S}_t$.

**Local parameter-information approximation.** Louis' identity [1] needs both moments, and we approximate each term of (S7). In the complete-information term we drop the second-derivative dynamics term $\nabla^2_{\boldsymbol{\theta}} \hat{\mathbf{z}}_t^-$, a Gauss–Newton approximation in $\boldsymbol{\theta}$ that is exact when $\hat{\mathbf{z}}_t^-$ is affine in $\boldsymbol{\theta}$ and otherwise discards a contribution proportional to the residual $\mathbf{z}_t - \hat{\mathbf{z}}_t^-$. In the missing-information term we approximate $\mathrm{Cov}_{p_t}(\mathbf{z}_t) \approx \mathbf{P}_{z,t}$. Together, these approximations give the positive parameter-information approximation $\mathbf{I}_{\theta,t} := -\mathbf{H}_{\theta,t}$ from the main text, where $\mathbf{H}_{\theta,t}$ approximates $\nabla^2_{\boldsymbol{\theta}} \ell_t$ at $\hat{\boldsymbol{\theta}}_{t-1}$:

$$\mathbf{I}_{\theta,t} := \mathbf{S}_t^\top \left[(\mathbf{P}_{z,t}^-)^{-1} - (\mathbf{P}_{z,t}^-)^{-1} \mathbf{P}_{z,t} (\mathbf{P}_{z,t}^-)^{-1}\right] \mathbf{S}_t \tag{S13}$$

The bracket is not a separate modeling choice: substituting $\mathbf{P}_{z,t}$ from (S11) and applying the identity $\mathbf{A} - \mathbf{A}(\mathbf{A}+\mathbf{B})^{-1}\mathbf{A} = \mathbf{B}(\mathbf{A}+\mathbf{B})^{-1}\mathbf{A}$ with $\mathbf{A} = (\mathbf{P}_{z,t}^-)^{-1}$ and $\mathbf{B} = \mathbf{I}_{z,t}$ collapses it to

$$(\mathbf{P}_{z,t}^-)^{-1} - (\mathbf{P}_{z,t}^-)^{-1} \mathbf{P}_{z,t} (\mathbf{P}_{z,t}^-)^{-1} = \mathbf{I}_{z,t} \left(\mathbf{I} + \mathbf{P}_{z,t}^- \mathbf{I}_{z,t}\right)^{-1}, \tag{S14}$$

so that (S13) is $\mathbf{I}_{\theta,t} = -\mathbf{H}_{\theta,t}$, the negative of the Hessian increment in (12). The same score and precision follow from the joint Gauss–Newton quadratic model of the log posterior in $(\delta\mathbf{z}_t, \delta\boldsymbol{\theta}_t)$ [2]: eliminating $\delta\mathbf{z}_t$ through the Schur complement of the state block, whose inverse is $\mathbf{P}_{z,t}$, gives the precision $\boldsymbol{\Lambda}_{\theta,t}^- - \mathbf{H}_{\theta,t}$ and the right-hand side $\mathbf{s}_{\theta,t}$ of (9) when $\mathbf{S}_t = \mathbf{F}_{\theta,t}$. This equivalence holds for the one-step parameter update only; it does not extend to the full dual filter or to the block sensitivity recursion (13).

**Role of the missing-information term.** Equation (S14) identifies the missing-information term as the source of the uncertainty discount discussed in Section II-C2. Retaining only the complete-information term would leave $\mathbf{S}_t^\top (\mathbf{P}_{z,t}^-)^{-1} \mathbf{S}_t$, which is the limit as all eigenvalues of $\mathbf{I}_{z,t}$ tend to infinity of (S14): it is the information the parameters would receive if the measurement resolved the latent state exactly, leaving $\mathbf{P}_{z,t} \to \mathbf{0}$. Since $\mathbf{I}_{z,t}(\mathbf{I}+\mathbf{P}^{-}_{z,t}\mathbf{I}_{z,t})^{-1} \preceq (\mathbf{P}^{-}_{z,t})^{-1}$, dropping the term never understates $\mathbf{I}_{\theta,t}$, and it credits the parameters with information that the observation in fact left in the latent state whenever the missing-information correction is nonzero. This complete-data limit differs from simply omitting the uncertainty discount, which gives $\mathbf{S}_t^\top \mathbf{I}_{z,t}\mathbf{S}_t$.

##### *A.3 Coarse-grid planning and adaptive intervals*

The planner of Section II-D uses two practical modifications that change the planning time grid and the replanning interval without changing the acquisition objective.

**Coarse-grid planning.** During iCEM search, we scale the model time step to $c\Delta t$, extending the horizon to $cH$ environment steps while retaining $H$ planning steps. We execute the first $n_c = \lceil \frac{K_u}{c} \rceil$ coarse inputs, each repeated for $c$ environment steps, and truncate the sequence to $K_u$ inputs.

Coarse rollouts use Euler steps of $c\Delta t$, $\tilde{\mathbf{F}}_z = \mathbf{I} + c\Delta t\, \partial\mathbf{v}/\partial\mathbf{z}$, $\tilde{\mathbf{F}}_\theta = c\Delta t\, \partial\mathbf{v}/\partial\boldsymbol{\theta}$, and $c$ times the per-step process-noise covariance. Each coarse transition spans $c$ observation bins but uses the state Fisher information $\tilde{\mathbf{I}}_{z,k}$ of a single bin. The accumulated information $-\sum_k \tilde{\mathbf{H}}_{\theta,k}$ in (21) can therefore be up to $c$ times smaller than the information that all $c$ bins provide, so the objective weighs predicted information less against the current precision $\boldsymbol{\Lambda}_{\theta,t}$.

**Adaptive update and replanning intervals.** The fixed-interval planner updates the parameter posterior every $K_\theta$ observations and replans every $K_u$ inputs. We also consider adaptive interval rules that preserve these intervals as maximum delays but allow the algorithm to update or replan earlier when the accumulated evidence or tracking error warrants it.

For parameter updates, let the block Hessian $\mathbf{H}_{\theta,\mathcal{T}_m}$ of (14) sum over the observations $t_m+1,\ldots,t$ received so far in the current block. With $\boldsymbol{\Lambda}_{\theta,t} = \mathbf{P}_{\theta,t}^{-1}$, we score this block by the posterior-normalized information gain

$$\begin{aligned}\mathcal{A}_{\theta,t} &= \frac{1}{2d_\theta}\log\det\Big(\mathbf{I} - \boldsymbol{\Lambda}_{\theta,t}^{-1}\mathbf{H}_{\theta,\mathcal{T}_m}\Big)\\ &= \frac{1}{2d_\theta}\log\det\Big(\mathbf{I} - \mathbf{P}_{\theta,t}^{1/2}\mathbf{H}_{\theta,\mathcal{T}_m}\mathbf{P}_{\theta,t}^{1/2}\Big)\end{aligned} \tag{S15}$$

where $d_\theta$ is the dimension of the inferred parameter vector. After a minimum delay $K_{\theta,\min}$, we apply the blockwise Gaussian parameter update as soon as

$$n_{\theta,t} \geq K_{\theta,\min} \quad \text{and} \quad \mathcal{A}_{\theta,t} \geq \kappa_\theta, \tag{S16}$$

where $n_{\theta,t}$ is the number of observations in the current block and $\kappa_\theta$ is the information-gain threshold. If this threshold is not met, the update is still applied at the maximum interval $n_{\theta,t} = K_\theta$. Whenever a parameter update is applied, the policy forces a replan because the predictive dynamics and information objective have changed.

For replanning based on model mismatch, we compare the filtered state after the new observation with the state predicted by the plan currently being executed. If $\hat{\mathbf{z}}_t$ is the filtered state mean and $\tilde{\mathbf{z}}_{t|t_0}$ is the planned state at the same execution index (with $t_0$ the time at which the current plan was computed), the normalized tracking error is

$$\mathcal{E}_{z,t} = \frac{1}{d_z}\left(\hat{\mathbf{z}}_t - \tilde{\mathbf{z}}_{t|t_0}\right)^\top \mathbf{P}_{z,t}^{-1}\left(\hat{\mathbf{z}}_t - \tilde{\mathbf{z}}_{t|t_0}\right), \tag{S17}$$

where $d_z$ is the state dimension and $\mathbf{P}_{z,t}$ is the current filtering covariance.

After a minimum execution delay $K_{u,\min}$, the planner replans early when

$$n_{u,t} \geq K_{u,\min} \quad \text{and} \quad \mathcal{E}_{z,t} > \kappa_z, \tag{S18}$$

with $\kappa_z$ the tracking-error threshold and $n_{u,t}$ the number of inputs executed from the current plan. This rule triggers replanning when the realized trajectory has deviated from the previously planned trajectory relative to the posterior state uncertainty. If no state-tracking or parameter-update trigger occurs, the planner replans at the nominal interval $K_u$.

### APPENDIX B
### EXPERIMENTAL DETAILS AND SUPPLEMENTARY RESULTS

#### B.1 Experimental settings

**Runs and evaluation.** Each benchmark condition uses 100 random seeds and 2000 environment interactions, except the objective-ablation system, which uses 750 interactions (Appendix B.4); the spiking-network experiment uses 20 seeds and 20 identification trials (Section III-E). Seeds control the initial state, process and observation noise, and initial parameter belief. For the global vector-field rollout score, we draw 100 initial states uniformly from the evaluation bounds and simulate the true and learned dynamics without control for 200 steps.

**Benchmark settings.** Wilson–Cowan uses $q_z = 0.1$, input bound 3, initial states drawn uniformly from $[-2, 2]^2$, which is the rate range $[0, \frac{1}{2}]^2$, and evaluation bounds $[-2.2, 2.2]^2$. The Wilson–Cowan weights have magnitudes of order 10 and are positive by definition, so each initial weight guess is $10|\xi_i|$ with $\xi_i \sim \mathcal{N}(0, 1)$, and the initial parameter covariance is $100\,\mathbf{I}$. All methods use the same guesses for a given seed. The Wilson–Cowan observation study changes only the loading calibration and geometry (Section III-C).

**Wilson–Cowan model.** The gain functions in (25) are $S_j(x) = 1/(1 + e^{-a_j(x-\vartheta_j)}) - 1/(1 + e^{a_j\vartheta_j})$ for $j \in \{E, I\}$. We use a standard parameter set of the original model with two stable states [3]: true weights $w_{EE} = 12$, $w_{EI} = 4$, $w_{IE} = 13$, and $w_{II} = 11$, and fixed gains $a_E = 1.2$ and $a_I = 1$, thresholds $\vartheta_E = 2.8$ and $\vartheta_I = 4$, and refractory constants $k_E = 0.97$ and $k_I = 0.98$. The refractory factors keep both rates below 0.49 without input, and the latent state $\mathbf{z}_t = 8([E_t, I_t]^\top - \frac{1}{4})$ maps the rate range $[0, \frac{1}{2}]$ to $[-2, 2]$. The circuit has a stable down state at $E = I = 0$, a stable up state at $(E, I) = (0.44, 0.23)$, and a saddle at $(0.19, 0.07)$. We simulate (26) with step size $\Delta t = 0.01$, giving per-step process-noise covariance $q_z \Delta t\,\mathbf{I}$. The transition is $f(\mathbf{z}, \mathbf{u}; \boldsymbol{\theta}) = \mathbf{z} + \Delta t[\mathbf{v}(\mathbf{z}; \boldsymbol{\theta}) + \mathbf{u}]$, so $\mathbf{F}_z = \mathbf{I} + \Delta t\, \partial\mathbf{v}/\partial\mathbf{z}$ and $\mathbf{F}_\theta = \Delta t\, \partial\mathbf{v}/\partial\boldsymbol{\theta}$.

**Policy settings.** On the Wilson–Cowan benchmark, the PALDI planner uses 20 planning transitions with coarse factor 20, a 400-step horizon, nominal intervals $K_\theta = K_u = 20$, minimum delays $K_{\theta,\min} = K_{u,\min} = 5$, and block-gain and tracking-error thresholds $\kappa_\theta = 0.001$ and $\kappa_z = 3$. We selected $\kappa_\theta$ among 16 settings on seeds 100–119, which are not among the reported runs. In the spiking-network experiment, PALDI instead plans 20 transitions with coarse factor 2, a 200-ms horizon within the 2-s trials, with minimum delays $K_{\theta,\min} = K_{u,\min} = 3$ and $\kappa_\theta = 0.05$. The settings of the objective-ablation planners are given in Appendix B.4. Every policy that uses the update of Section II-C2 sets $q_\theta = 5 \times 10^{-6}$ per observation in (8), which is $10^{-4}$ per nominal 20-observation block ($5 \times 10^{-4}$ per observation for the Wilson–Cowan weights, the same value on their scale of 10). Because the inflation scales with the number of observations, learners that update at every step, such as Myopic, PRBS, and random inputs, accumulate the same inflation as learners with 20-observation blocks. iCEM uses 24 candidate sequences, 4 iterations, 6 elites, initial standard deviation 0.5, mean smoothing 0.1, colored sampling noise with exponent 1, and 30% elite reuse. Parameter means are clipped to $[-5s, 5s]$, with $s = 10$ for the Wilson–Cowan weights and $s = 1$ otherwise, and $10^{-6}\mathbf{I}$ is added to the predictive state covariance. The agents' filters use the process-noise covariance $\log(1 + q_z)\,\Delta t\,\mathbf{I}$, which is 4.7% below the simulator's $q_z \Delta t$ for Wilson–Cowan.

**Observation calibration.** For the benchmark readouts, the Fisher SNR is

$$10 \log_{10} \frac{\sum_i \langle x_i^2 \rangle}{\left\langle \operatorname{tr}\left[(\mathbf{C}^\top \operatorname{diag}(\boldsymbol{\lambda}(\mathbf{x}))\mathbf{C} + 10^{-6}\mathbf{I})^{-1}\right]\right\rangle},$$

where $\mathbf{x}$ is the latent state standardized over 100 input-free, noise-free 500-step trajectories, $\boldsymbol{\lambda}(\mathbf{x})$ is the count mean per bin, and $\langle\cdot\rangle$ averages over the trajectory states [4]. NeuroFisherSNR scales $\mathbf{C}$ and sets the bias to reach the target under the mean- and maximum-rate targets. The uniform Wilson–Cowan readout spaces the loading directions every 18°, with per-neuron gains drawn once and shared across runs. The biased readout keeps the first (excitatory) loading component at its drawn scale, makes the second (inhibitory) component positive and twice as large, and points one row at 45°, the only row whose rate rises with the excitatory state; the calibration then reaches $-6$ dB against the $-5$ dB target. For the Wilson–Cowan readout, one model time unit is taken as 1 s, so each step is a 10-ms bin.

#### B.2 Exploration and parameter information in Wilson–Cowan

Figure S1 compares the saved trajectories from the Wilson–Cowan condition with uniform loading at $-5$ dB in Figure 3A. Each method has 100 complete runs. We divide $[-7.5, 7.5]^2$, which contains all recorded states, into $100 \times 100$ cells of width 0.15 in each latent coordinate. This domain extends beyond the rate range because the additive input can move the state to $z_i < -2$, which corresponds to a negative rate. Coverage is the percentage of cells containing at least one recorded state in a run. It measures sampled coverage at this resolution; we do not fill the area between trajectory points. With $40 \times 40$ or $80 \times 80$ grids, PALDI keeps the highest median final coverage.

For a common parameter-information diagnostic, we evaluate the local Jacobians at the true weights and recorded states, and the Poisson Fisher information at each recorded successor state using the saved readout. We propagate state covariance and measurement-corrected sensitivity as in (4), (5), and (13), using one block over the full trajectory, initial state covariance $\mathbf{I}_2$, and zero initial sensitivity. Define

$$\mathcal{G}_T = \frac{1}{2}\log\det\left(\mathbf{I}_4 + \mathbf{P}_{\theta,0}\sum_{t=1}^{T}\mathbf{S}_t^\top(\mathbf{I}_{z,t}^{-1} + \mathbf{P}_{z,t}^{-})^{-1}\mathbf{S}_t\right), \qquad \mathbf{P}_{\theta,0} = 100\,\mathbf{I}_4 \tag{S19}$$

We use the same calculation for every method, without parameter diffusion, sensitivity resets, or shrinkage.

PALDI has the highest median final coverage (2.40%) and information proxy (11.27 nats). FLEX covers fewer cells than RHC-US but reaches a larger information proxy, so coverage alone does not determine how informative the visited states are. These diagnostics support the exploration interpretation of Section III-B, without establishing a causal explanation of the recovery differences.

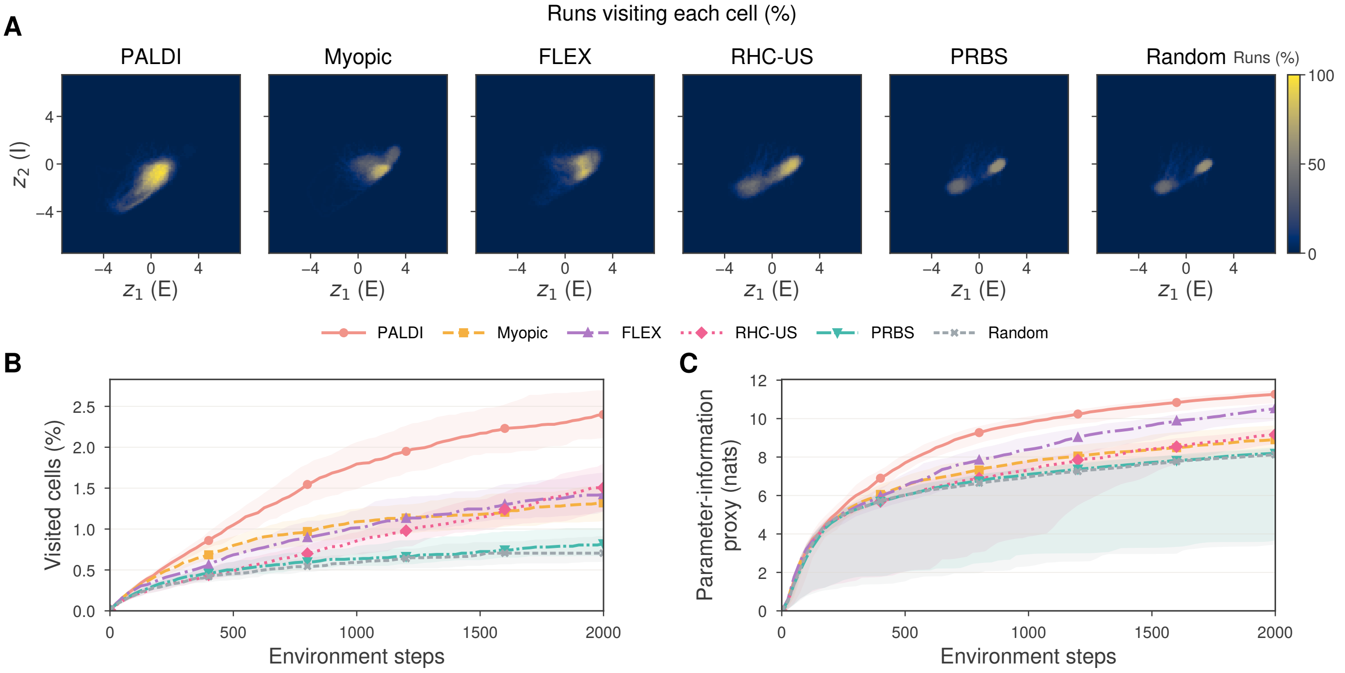


Fig. S1. **Exploration in the neural mass model.** State-space coverage and parameter information by method. (**A**) Percentage of runs that visit each cell. (**B**) Cumulative fraction of cells visited per run. (**C**) Information proxy of (S19), shown every 25 steps and at the last step. FLEX uses the stabilized update with filtered states, as in the main comparison. Medians and IQR across 100 seeds.

##### *B.3 FLEX state source and update variants*

The FLEX baseline of Section III-A1 uses the filtered state $\hat{\mathbf{z}}_t$ with the stabilized parameter update. This section separates the two adaptations by comparing three variants that share the acquisition objective (23), the interaction budget, and the environment presets, and differ only in the state supplied to the update and in whether the acceptance test is applied: the true state $\mathbf{z}_t$ with the original parameter update, the filtered state with the original parameter update, and the filtered state with the stabilized update (the baseline of the main comparison).

We accept an update only if the new parameters are finite, each lies in $[-5s, 5s]$, and $\|\Delta\hat{\boldsymbol{\theta}}_t\|_2 \leq 0.25\,s$, where the parameter scale is $s = 10$ for the Wilson–Cowan weights and $s = 1$ otherwise; the Gram-matrix regularization is divided by $s^2$. Otherwise, both the parameters and Gram matrix remain unchanged.

With uniform loading at $-5$ dB, the true-state variant reaches a final median $R^2_{\text{VF-roll}}$ of 0.56 (IQR $-0.73$ to 0.89). The filtered variants reach $-1.98$ (IQR $-2.40$ to $-1.48$) with the original update and 0.835 (IQR 0.79–0.87) with the stabilized update (Figure S2A). The true-state variant does not use the observations, but the sampled environment noise differs between conditions, so its median varies across panels: 0.66 under biased loading, and 0.58 and 0.08 at $-10$ and $-15$ dB. With the filtered state, the stabilized variant reaches 0.825 under biased loading and 0.765 and 0.568 at $-10$ and $-15$ dB (Figure S2B–D).

FLEX regresses the finite-difference target $(\hat{\mathbf{z}}_{t+1} - \hat{\mathbf{z}}_t)/\Delta t$. The EKF predicts with the current FLEX estimate, so this target contains the filter correction divided by $\Delta t$. The filter starts from a fixed mean with unit covariance, so the first corrections are large. With a small Gram-matrix regularization, the first update nearly fits this outlier, and with the original update the estimates diverge after it. The stabilized variant rejects such updates and keeps the previous parameters and Gram matrix.

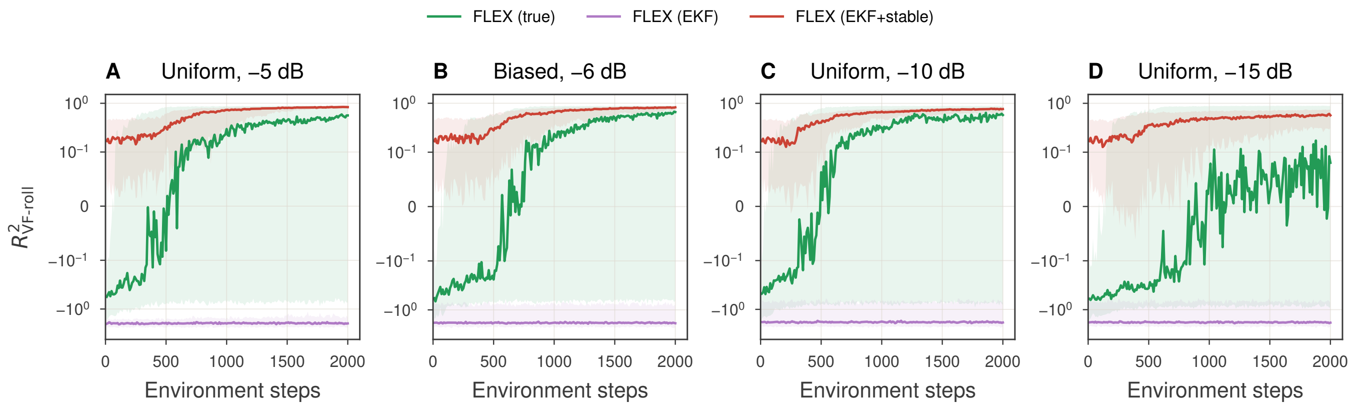


Fig. S2. **Comparison of FLEX variants.** $R^2_{\text{VF-roll}}$ of FLEX on the neural mass model with the true state and the original update (true), the filtered state and the original update (EKF), and the filtered state and the stabilized update (EKF+stable). (**A**) Uniform loading at $-5$ dB. (**B**) Biased loading at $-6$ dB. (**C,D**) Uniform loading at $-10$ and $-15$ dB. Medians and IQR across 100 seeds, on a symmetric logarithmic axis.

##### *B.4 Construction of the four-region system*

We set $\boldsymbol{\theta}^{\text{true}} = (1, 1, 1)$ and initialize all three parameter means at zero with covariance $\mathbf{I}$. The true state and the filter mean start at $(r, \mathbf{s}, \mathbf{b}, h) = (-1, \mathbf{0}, \mathbf{0}, 0)$, with filter covariance $\mathbf{I}$. The active objectives use the adaptive update and replanning rules of Appendix A.3. We run each method for 750 interactions across 100 matched seeds.

The region functions $g_j(r) = \exp[-(r-c_j)^2/(2w^2)]$, with $j \in \{D, N, B, F\}$, use centers $(c_D, c_N, c_B, c_F) = (-1.8, -0.5, -0.1, 0.3)$ and width $w = 0.1$. The passive equilibrium $r = -1$ lies outside the regions, with region D on one side and regions N, B, and F on the other. Without the domain boundary, holding D, N, B, and F requires inputs $(-0.8, 0.5, 0.9, 1.3)$; the acquisition objective has no input cost. The paired positively and negatively tuned Poisson neurons have loading magnitudes $\mathbf{a} = (a_1, \ldots, a_8) = (0.3, 0.3, 0.3, 0.3, 0.03, 0.03, 0.03, 0.05)$ in $\mathbf{C}$ for the coordinates $(r, \mathbf{s}, \mathbf{b}, h)$, a 50 Hz baseline per neuron and $\Delta t = 0.01$ s bins.

**Domain boundary.** To keep simulated states within a prescribed domain, we combine a smooth inward drift with projection onto the ball $\mathcal{B}_R = \{\mathbf{z} : \|\mathbf{z}\|_2 \le R\}$.

**Objectives.** All active policies share inference, learning, the planning time grid, and optimization; only the acquisition objective changes (Table I). The State Information objective removes parameter sensitivity. At each planning step $k$, it scores the reduction in the log-determinant of the state covariance by one observation,

$$\log\det \tilde{\mathbf{P}}^{-}_{z,k} - \log\det\left((\tilde{\mathbf{P}}^{-}_{z,k})^{-1} - \tilde{\mathbf{H}}_{z,k}\right)^{-1} = \log\det\left(\mathbf{I} - \tilde{\mathbf{P}}^{-}_{z,k}\tilde{\mathbf{H}}_{z,k}\right), \tag{S20}$$

where $((\tilde{\mathbf{P}}^{-}_{z,k})^{-1} - \tilde{\mathbf{H}}_{z,k})^{-1}$ is the state covariance after a Laplace update, and sums this score over the horizon. It therefore favors states that the observations inform, regardless of whether the parameters affect the dynamics there. The Perfect-observation objective removes the state Fisher information. It replaces the per-step parameter information $-\tilde{\mathbf{H}}_{\theta,k} = \tilde{\mathbf{S}}_k^{\top}(\tilde{\mathbf{I}}_{z,k}^{-1} + \tilde{\mathbf{P}}^{-}_{z,k})^{-1}\tilde{\mathbf{S}}_k$ ((S14)) with its limit for $\tilde{\mathbf{I}}_{z,k} \to \infty$, the upper bound $\tilde{\mathbf{S}}_k^{\top}(\tilde{\mathbf{P}}^{-}_{z,k})^{-1}\tilde{\mathbf{S}}_k$, and scores $\log\det\bigl(\mathbf{I} + \mathbf{P}_{\theta,t}\sum_{k=1}^{H}\tilde{\mathbf{S}}_k^{\top}(\tilde{\mathbf{P}}^{-}_{z,k})^{-1}\tilde{\mathbf{S}}_k\bigr)$. It keeps the parameter sensitivity and the uncertainty discount, treats every state coordinate as perfectly observed, and does not depend on the observation model. Both objectives use the same prediction-only rollouts of $\tilde{\mathbf{P}}^{-}_{z,k}$ and $\tilde{\mathbf{S}}_k$ as p-EIG.

The two disagreement objectives draw 8 parameter samples $\boldsymbol{\theta} \sim q_t = \mathcal{N}(\hat{\boldsymbol{\theta}}_t, \mathbf{P}_{\theta,t})$ at each evaluation and use the same samples for every candidate input sequence. For each sample, the learned dynamics are rolled out from the filtered mean $\hat{\mathbf{z}}_t$ under the candidate sequence without process noise. Disagreement in observations sums $\log(1 + v)$ over neurons and planning steps, where $v$ is the sample variance of a neuron's predicted count mean $\tilde{\lambda}_{j,k}$; disagreement in state sums the trace of the sample covariance of the predicted states $\tilde{\mathbf{z}}_k$ over planning steps. Neither objective accounts for latent-state uncertainty.

**Evaluation.** For this system, $R^2_{\text{VF-roll}}$ uses 100 initial states drawn uniformly from $r \in [-1.05, 0.35]$, $s_i \in [-0.25, 0.25]$, and $b_i \in [-0.25, 0.25]$, with $h = 0$. The true and learned dynamics are rolled out for 200 steps without input, noise, or boundary, and the score averages the $R^2$ of $s_1$, $s_2$, and $s_3$ rather than pooling all latent coordinates. The true-model reference is therefore exactly one. Region D lies outside the evaluation range of $r$, but $\mathbf{b}$ and $\mathbf{s}$ share $\boldsymbol{\theta}$, so data from region D identify the scored dynamics when the observations resolve $\mathbf{b}$.

##### *B.5 Spiking-network experimental protocol*

**Network and recorded state.** The data generator is the decision network of Wang [5], simulated in Brian 2 [6], with 1600 excitatory and 400 inhibitory leaky integrate-and-fire neurons. Two selective excitatory pools contain 240 neurons each and compete through shared inhibition. We record 20 neurons from each selective pool in 5-ms bins, giving $d_y = 40$. The two inputs inject current into the selective pools, with one input unit equal to 20 pA. We define the latent proxy $\mathbf{z}_t = 4(\mathbf{s}_t - \frac{1}{2}\mathbf{1})$, where $s_{i,t}$ is the pool-averaged NMDA gating variable. This proxy summarizes the network state; it is not a closed two-dimensional description of the spiking dynamics.

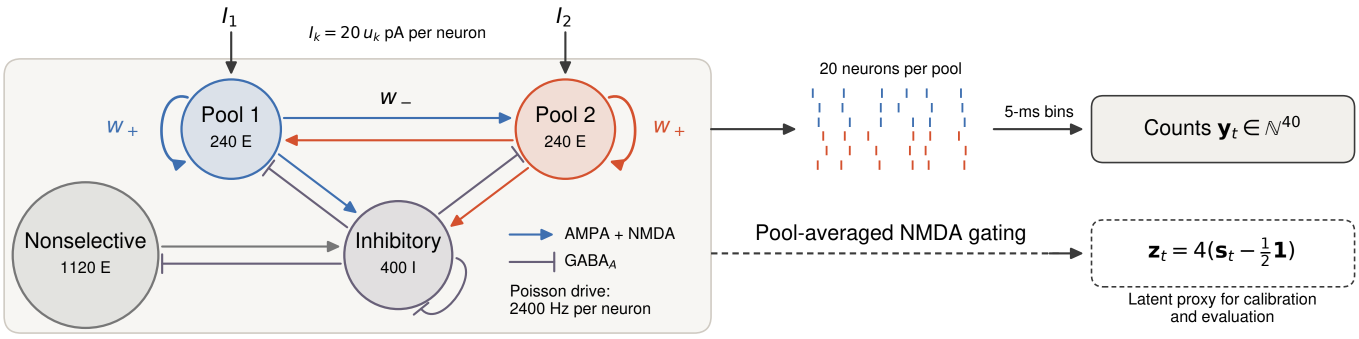


Fig. S3. **Wang spiking circuit.** Population structure and observations of the circuit [5], which motivates the Wong–Wang reduction [7]. Two selective pools of 240 excitatory neurons each, 1120 nonselective excitatory neurons, and 400 inhibitory neurons; arrows denote AMPA/NMDA excitation and bars GABA$_A$ inhibition. The network couplings, $w_+ = 1.7$ within a selective pool and $w_- = (1 - 0.15w_+)/0.85 \approx 0.876$ across pools, are not the fitted reduced-model parameters. Excitatory connections of the nonselective population are omitted except its input to the inhibitory population. Every neuron receives independent 2400-Hz Poisson background drive, and one input unit injects 20 pA into each neuron of the selected pool. We record 20 neurons per selective pool in 5-ms bins (40 channels; spike marks are schematic). The latent proxy, NMDA gating averaged over each pool, is used for readout calibration and model evaluation.

**Observation calibration.** The agent uses a Poisson log-linear count model, $\boldsymbol{\lambda}_t = \Delta t \exp(\mathbf{C}\mathbf{z}_t + \mathbf{b}_r)$, with $\mathbf{C} \in \mathbb{R}^{40\times 2}$. Time in the reduced model is measured in units of the 100-ms NMDA time constant, so $\Delta t = 0.05$ corresponds to one 5-ms bin. The bias uses these normalized time units. For each training seed, we calibrate $\mathbf{C}$ and $\mathbf{b}_r$ on 800 bins of open-loop drive and then hold them fixed. This calibration uses the network's recorded latent proxy as well as spike counts. Subsequent online inference and learning use spikes and known trial events. Thus the experiment assumes a calibrated observation map; it does not learn the latent coordinates from spikes alone. The observed counts come from the network, rather than from this fitted Poisson model.

**Identification trials.** We compare PALDI, Myopic, FLEX, RHC-US, PRBS, and random inputs using the baseline definitions in Section III-A1. Each policy receives 20 identification trials for each of 20 matched training seeds. A trial starts at $\mathbf{z} = (-2, -2)^\top$, and the network makes a decision when $|z_1 - z_2| > 2$. In trials 1–5, the network also receives the task's evidence, 6 pA to a randomly selected pool during the first second, and resets 200 ms after the decision, subject to a 2-s trial limit. Trials 6–20 have no evidence and always last 2 s; a decision does not end them. Agents know the evidence (pool and timing) and the trial rule but learn of a reset only when it occurs: the state filter then resets to the known state with covariance $0.01\mathbf{I}$. Planning rollouts do not simulate decisions or resets; a rollout continues the current trial for its whole horizon. Policies share the readout calibration and initialization for a given seed. As a passive reference, a learner with the per-step update of random inputs receives evidence in all 20 trials and applies no input; each of its trials ends 200 ms after the decision or at the 2-s limit. The duration of trials 1–5 depends on the selected inputs, so equal trial counts do not imply equal numbers of observed bins. The policies constrain inputs to the ball $\|\mathbf{u}_t\|_2 \le u_{\max}$, where $u_{\max} = 0.5$ (10 pA) in trials 6–20 and $u_{\max} = 0.1$ (2 pA) in trials 1–5. The reduced bound keeps each agent's input below the evidence: across the six policies, 597 of the 598 decisions in trials 1–5 match a replay with the same network noise and no input, which always chooses the evidence pool. One coarse planning step of PALDI lasts 10 ms, with 20 steps giving a 200-ms horizon. The Euler step of 0.1 in normalized time is locally stable near the decided fixed points of the fitted reduced model (multipliers 0.785 and 0.923).

**Prediction metric.** After each trial, we evaluate the estimated model on 12 held-out network trajectories under piecewise-constant random drives. Each 100-bin rollout starts from the recorded network latent proxy, with starts spaced by 20 bins, and uses the recorded inputs. We compute $R^2_{\text{roll}} = 1 - \text{SSE}/\text{SST}$ by pooling both coordinates, times, rollout starts, and trajectories; SST is measured about the pooled target mean. The score tests 500-ms prediction conditional on the correct initial latent state. It does not include filtering error and differs from $R^2_{\text{VF-roll}}$, which requires a known latent vector field. We report medians and IQRs across training seeds.

**Control task and references.** We freeze the learned parameters after $k \in \{0, 1, 5, 10, 15, 20\}$ trials and use the same task controller for every identification policy. Each task trial applies 6 pA of evidence to a randomly selected pool for 1 s. In the *change-of-mind* task, the network makes its initial decision without control. Control then lasts at most 2 s, and success means that the alternative pool leads the chosen pool, $z_a - z_c > 2$, within 2.5 s of the initial decision. Decision and success events are defined using the network latent proxy. The evidence and the alternative pool are known to the controller.

Physical control currents obey $\|\mathbf{i}_t\|_2 \le 20$ pA, with $\mathbf{i}_t = 20\,\mathbf{u}_t$ pA. We report the integrated squared-current budget $B = \Delta t_{\text{phys}} \sum_t \|\mathbf{i}_t\|_2^2$, where $\Delta t_{\text{phys}} = 0.005$ s. Because one model time unit is 0.1 s, $B = 40B_u$ pA$^2$ s for the implementation's normalized budget $B_u = 0.05 \sum_t \|\mathbf{u}_t\|_2^2$. We selected $B \in \{400, 480\}$ pA$^2$ s using calibration trials with the fixed open-loop inputs and separate seeds; the main text reports 480 pA$^2$ s, and Appendix B.7 reports 200, 400, 480, and 600 pA$^2$ s. Evidence is excluded. The budget sums the squared per-neuron currents across the two selective pools; it is not total circuit current or physical electrical energy. The controller filters spikes with the frozen model and replans using iCEM [8] every 200 ms. It scores a smooth Monte Carlo approximation to the probability of the task's first-passage success rule, subject to the remaining budget. The main control figure uses warm starts: each search includes simple pushes and the remaining previous plan. The task controller's iCEM samples 64 candidates, reduced by a factor of 1.25 per iteration to at least 16, with 8 elites, 8 iterations, initial standard deviation 0.5, and coarse factor 10 (50-ms holds). It plans over the rest of the control window plus 100 bins and scores the mean of $\sigma(4(G-2))$, where $G$ is the largest value of $z_a - z_c$, over 8 Monte Carlo rollouts with process variance 0.1 and common random numbers. Inputs are projected to $\|\mathbf{u}_t\|_2 \leq 1$ and to the remaining budget.

References use no input, Constant-amplitude input across the control window, Maximum-amplitude input from onset until the budget is spent, and the same iCEM controller with either the initial estimate or the privileged fitted model. Both fixed open-loop inputs excite the alternative pool and suppress the chosen pool along a unit-norm direction. The fitted-model controller still filters spikes using each seed's calibrated readout; it does not receive the online latent state. We use eight fixed network seeds, shared across controllers. One change-of-mind trial has no initial decision within 2 s and is excluded for every controller, leaving seven valid change-of-mind trials. Success is averaged over valid task trials and then training seeds. Error bars and bands show SEM across the 20 training seeds, conditional on this shared task set. The fixed open-loop inputs have one outcome per task seed and no across-training-seed error bar.

##### *B.6 Reduced-model parameterization and reference fit*

The reduced model follows Wong and Wang [7], who derive it from the spiking circuit [5]. We measure time in units of the NMDA time constant $\tau_N = 100$ ms and keep the shape of their effective firing response, $\phi(q) = q/(1 - e^{-q})$, with the fixed drive scale 6 in (28). The learned scale $\gamma$ absorbs the activation amplitude and the time normalization. The five learned coordinates are $\boldsymbol{\theta} = (w_+, w_-, h_{\text{raw}}, \gamma_{\text{raw}}, g_{\text{raw}})^\top$, with $h = h_{\text{raw}}/6$, $\gamma = \text{softplus}(\gamma_{\text{raw}})$, and $g = \text{softplus}(g_{\text{raw}})$, so $\gamma, g > 0$ while $w_+$ and $w_-$ are unconstrained. The mean transition is $f(\mathbf{z}, \mathbf{u}; \boldsymbol{\theta}) = \mathbf{z} + \Delta t\, \mathbf{v}(\mathbf{z}, \mathbf{u}; \boldsymbol{\theta})$ with $\Delta t = 0.05$, one 5-ms bin. Input to one pool reaches the other pool only through later changes in its gate, so the model omits faster cross-population effects through firing rates and shared inhibition. The coefficients are fitted to trajectories, so they are effective parameters that depend on the sampled operating regime.

Each identification policy learns all five coordinates online from its own spike observations and inputs. Initial parameter means are drawn from $\mathcal{N}((0, 0, 0, -0.661183, 0)^\top, \mathbf{I})$ across seeds, and each parameter belief starts with covariance $\mathbf{I}$. There is no exact $\boldsymbol{\theta}^{\text{true}}$ of the spiking network in this class; the fitted reference is neither biological ground truth nor an upper bound on prediction or control.

The fitted reference uses latent proxies from 24 trajectories with piecewise-constant inputs and 24 with task-like inputs. The latter combine 6-pA evidence for 1 s with 200-ms signed pushes in alternating 200-ms blocks. We minimize 100-bin rollout squared error from starts spaced by 20 bins, using float64 Adam for 300 iterations at learning rate 0.05 from 16 initializations. The fit is selected by training error alone. These data are separate from the identification trials and the 12 held-out prediction trajectories. The fitted reference reaches $R^2_{\text{roll}} = 0.969$ on those held-out trajectories.

##### *B.7 Control budget and success in the spiking network*

Figure S4 extends the change-of-mind comparison to budgets $B \in \{200, 400, 480, 600\}$ pA$^2$ s after 20 identification trials, using the same learned models, task trials, and warm-start controller as Figure 6.

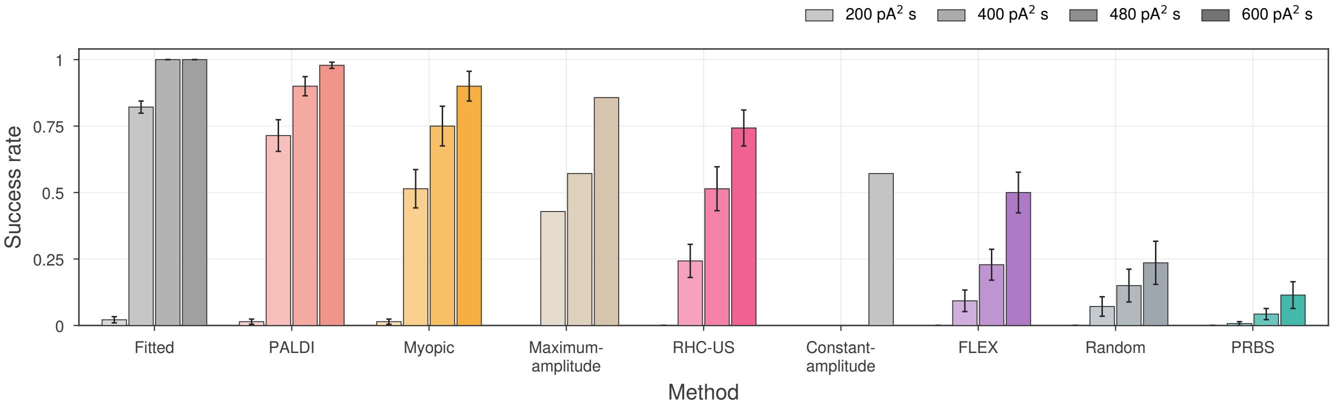


Fig. S4. **Control success across input budgets.** Change-of-mind success at budgets $B = \int \|\mathbf{i}_t\|^2 \, \mathrm{d}t \in \{200, 400, 480, 600\}$ pA$^2$ s. Methods are sorted by success at the highest budget; bars within a group follow increasing budget. Learned models are frozen after 20 identification trials and used with the same controller. Fitted uses one fitted model with the 20 seed-specific observation maps, so its error bars reflect only these maps. Constant- and Maximum-amplitude are fixed open-loop inputs and have no error bars. Zero-height bars denote zero success. Means and SEM across 20 training seeds, each evaluated on the same seven valid trials (one of eight trials has no initial decision).

##### *B.8 Median and interquartile-range summaries*

For the bistable benchmark, Section III reports the mean $R^2_{\text{VF-roll}}$ across the 100 seeds with the standard error of the mean. Large rollout errors can produce a long negative tail in the across-seed $R^2_{\text{VF-roll}}$ distribution. The mean is sensitive to this tail, whereas the median describes a typical run. A negative score alone does not establish numerical divergence. Figure S5 therefore repeats those comparisons using the median across seeds with the 25th–75th percentile interquartile range, computed over the same runs. For PALDI, Myopic, RHC-US, and the non-adaptive inputs, the order is the same as by the mean, apart from ties. The median of FLEX is above its mean in every condition: by the median, FLEX is above Myopic at $-10$ dB and above PALDI under biased loading, whereas by the mean it is below both.

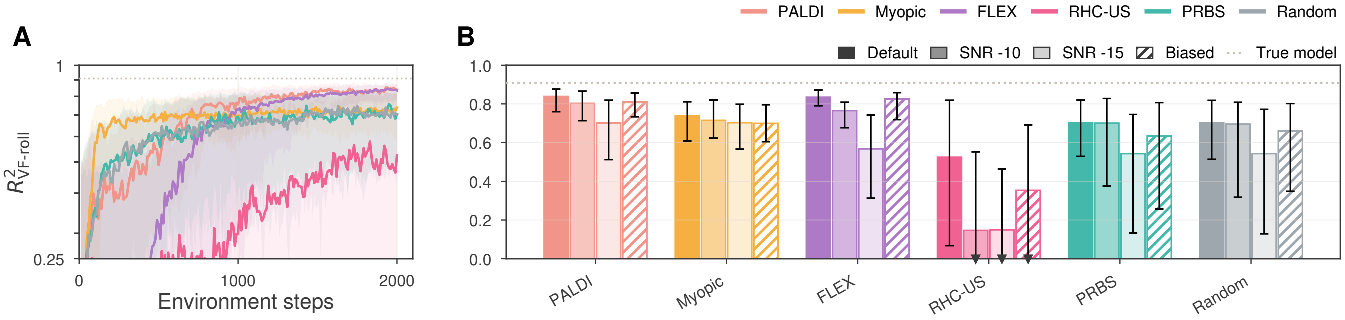


Fig. S5. **Observation conditions, median and IQR.** Median/IQR version of Figure 3.
